# A Unified Framework to Analyze Transposable Element Insertion Polymorphisms using Graph Genomes

**DOI:** 10.1101/2023.09.11.557209

**Authors:** Cristian Groza, Xun Chen, Travis J. Wheeler, Guillaume Bourque, Clément Goubert

**Affiliations:** Quantitative Life Sciences, McGill University, Montréal, QC, Canada; Institute for the Advanced Study of Human Biology (ASHBi), Kyoto University, Kyoto, Japan; R. Ken Coit College of Pharmacy, University of Arizona, Tucson, AZ, USA; Canadian Centre for Computational Genomics, McGill University, Montréal, QC, Canada; Victor Phillip Dahdaleh Institute of Genomic Medicine at McGill University, Montréal, QC, Canada; Human Genetics, McGill University, Montréal, QC, Canada

**Author notes:** R. Ken Coit College of Pharmacy, University of Arizona, Tucson, AZ, USA.

**Keywords:** Transposable elements, insertion polymorphism, graph genomes, structural variation, genotyping, genome evolution

## Abstract

Transposable Elements are ubiquitous mobile DNA sequences evolving among their hosts’ genomes, generating insertion polymorphisms that contribute to genomic diversity. We present GraffiTE, a flexible pipeline to analyze polymorphic mobile elements. By integrating state-of-the-art structural variant detection algorithms and graph genomes, GraffiTE identifies polymorphic mobile elements from genomic assemblies and/or long-read sequencing data, and genotypes these variants using short or long read sets. Benchmarking on simulated and real datasets reports high precision and recall rates. GraffiTE is designed to allow non-expert users to perform comprehensive analyses, including in models with limited transposable element knowledge and is compatible with various sequencing technologies. GraffiTE is freely available at https://github.com/cgroza/GraffiTE. Here, we demonstrate the versatility of GraffiTE by analyzing human, *Drosophila melanogaster,* maize, and *Cannabis sativa* pangenome data. These analyses reveal the landscapes of polymorphic mobile elements and their frequency variations across individuals, strains, and cultivars.

## Introduction

Transposable Elements (TEs) represent a heterogeneous collection of repetitive sequences found in virtually all eukaryotic species^1,2^. A unifying characteristic of TEs is their initial ability to multiply and colonize new loci within their host genome through transposition^3^. Active TE families (sets of copies descending from the same source sequence) are often quickly silenced through the rapid evolution of host defense mechanisms^4^. However, transposition of evolutionarily young TE families acts as a major source of structural variants (SVs) in a multitude of species^5^. As an example, and with only three TE families currently active, the human population harbors tens of thousands of polymorphic loci differentially present or absent between individuals^6,7^ (hereinafter referred to as polymorphic mobile elements, or pMEs). This number is even higher in *Drosophila melanogaster*, whose genome hosts many fewer TEs than the human genome (∼20 and ∼50%, respectively) but possesses a significantly higher proportion of families remaining active^8^.

With the advancement of sequencing technologies, and fostered by the intricate involvement of TEs with a wide array of biological processes^2^, the study of pMEs has risen in popularity. Indeed, as segregating variants, pMEs can be used as genetic markers, either to describe the population structure or to search for selection signatures^9–13^. Furthermore, the mutagenic effect of new insertions, coupled with the dissemination of their own regulatory and coding sequences among genomes, has been shown to affect regulatory networks and hosts’ phenotypes from diseases to adaptation^14–18^. In effect, pMEs are a significant contributor to genomic variation, constantly delivering raw material for evolution.

The detection and genotyping of pMEs has proven to be a challenging task, due to their repetitive nature and variable size (from hundreds to tens of thousands of bp)^19^. Nevertheless, a plethora of methods has been published, in particular targeting short-read paired-end datasets^6,20–27^. With the increased availability of long-read sequencing, new tools have emerged to detect and genotype SVs, including pMEs, with increased sensitivity^28–32^. Indeed, long reads are able to capture full-length pME sequences in context, i.e. with enough flanking sequence to anchor them with confidence in a reference genome. With long-read sequencing growing in popularity and affordability, an increasing number of projects are producing multiple high-quality, chromosome-level assemblies for a given species, referred to as pangenomes^33–37^.

Pangenomics, the integrative analysis of multiple related genomes simultaneously, has become the gold-standard in comparative genomics^38^. This new paradigm allows for a better representation of the genetic diversity in a given model, by including genomic variants previously ignored or missed, due to a lack of information stored in a single reference genome. As a consequence, pangenomes can dramatically enhance the estimation of SVs’ frequencies, which is for example extremely relevant in the context of rare diseases diagnostis^39^, agriculture^40,41^, and association studies in general^42^.

Among the tools developed for pangenome analysis, graph genomes are flexible data structures enabling representation of genomic variation, from single nucleotide polymorphisms (SNVs) to large SVs, in a single graph. In graph genomes, variants are represented by bubbles of divergent sequences (or nucleotides) and shared segments are collapsed into a single node^43^. Graph genomes allow high-precision mapping of genomic, transcriptomic^44^ and epigenomic^45,46^ variants, enabling research in evolutionary and functional genomics.

Existing pME methods that can apply to pangenome projects may be restricted to the detection of non-reference variants (i.e. TE absent from a reference genome, for example TLDR and TELR^28,29^), or specialized to a specific type of TE or organism^31^. Furthermore, direct detection of pMEs from alternative genomic assemblies is rare (though this feature – restricted to a single alternative assembly at a time – exists in TrEMOLO^30^). Finally, to our knowledge, none of the existing methods offer a graph genome framework dedicated to detect and genotype pMEs in a reliable yet flexible way.

To fill this gap, we have created GraffiTE (pronounced “gruh·FEE·tee”, like the popular street art form, *graffiti*). GraffiTE is a general-purpose pipeline for calling and genotyping pMEs, applicable to any model for which a list of consensus TE sequences of interest is available. GraffiTE makes available different state-of-the-art methods to detect and report pMEs from genomic assemblies or long-reads, and performs graph genotyping using short or long read data. The GraffiTE workflow is managed using the Nextflow framework^47^. GraffiTE was built with non-expert users in mind: a complete analysis of multiple genomic samples can be performed in a single command, while dependencies are directly available through containerization. Furthermore, variant handling, comparison and sharing is facilitated through the use of standardized variant call format (VCF). Finally, advanced configuration and optimisation are available for high performance clusters or cloud deployment.

After a brief presentation of the method, we will present the results of a synthetic and real data benchmark to demonstrate the performance of the tool. Following that, we show the versatility of the tool by searching for pMEs using 47 diploid human genome assemblies, 30 haploid *Drosophila melanogaster* assemblies alongside their original long-read sets, 23 long read set of various maize cultivars to annotate polymorphisms at the *bz* locus. Finally, we demonstrate how the tool can be applied to a species with limited knowledge about TEs, the emerging agricultural model *Cannabis sativa*.

## Results

### A unified framework to study TE insertion polymorphisms

GraffiTE organizes a complex collection of SVs and pangenome analysis software in order to identify, extract, and analyze genomic variants likely to represent mobile element insertion polymorphisms (pMEs). The pipeline is implemented in the Nextflow framework, and relies on an Apptainer container (formerly known as Singularity^48^) to provide all the required dependencies.

In brief, the pipeline is divided into three main steps (Figure 1). First, SVs are searched between either (i) one or more alternative assemblies and a reference genome or (ii) long-read sets and a reference genome (both searches can be also combined). Second, SVs found in individual samples are merged and further filtered to retain pMEs. Alternatively, a user-provided VCF file reporting both reference and alternative alleles sequences for each variant can be used as input. Finally, each pME detected can be further genotyped by mapping short or long reads against a TE graph genome. This graph genome represents each identified pME as a bubble, i.e. providing alternate paths in the graph, where both presence and absence alleles are available for read mapping and genotyping. The first two steps focus on pME detection, i.e. establishing whether a given TE copy is present or absent in a sample. The third step, called pME genotyping, is aimed to report whether each pME detected is homozygous or heterozygous (provides allele frequencies).

**Figure 1.**
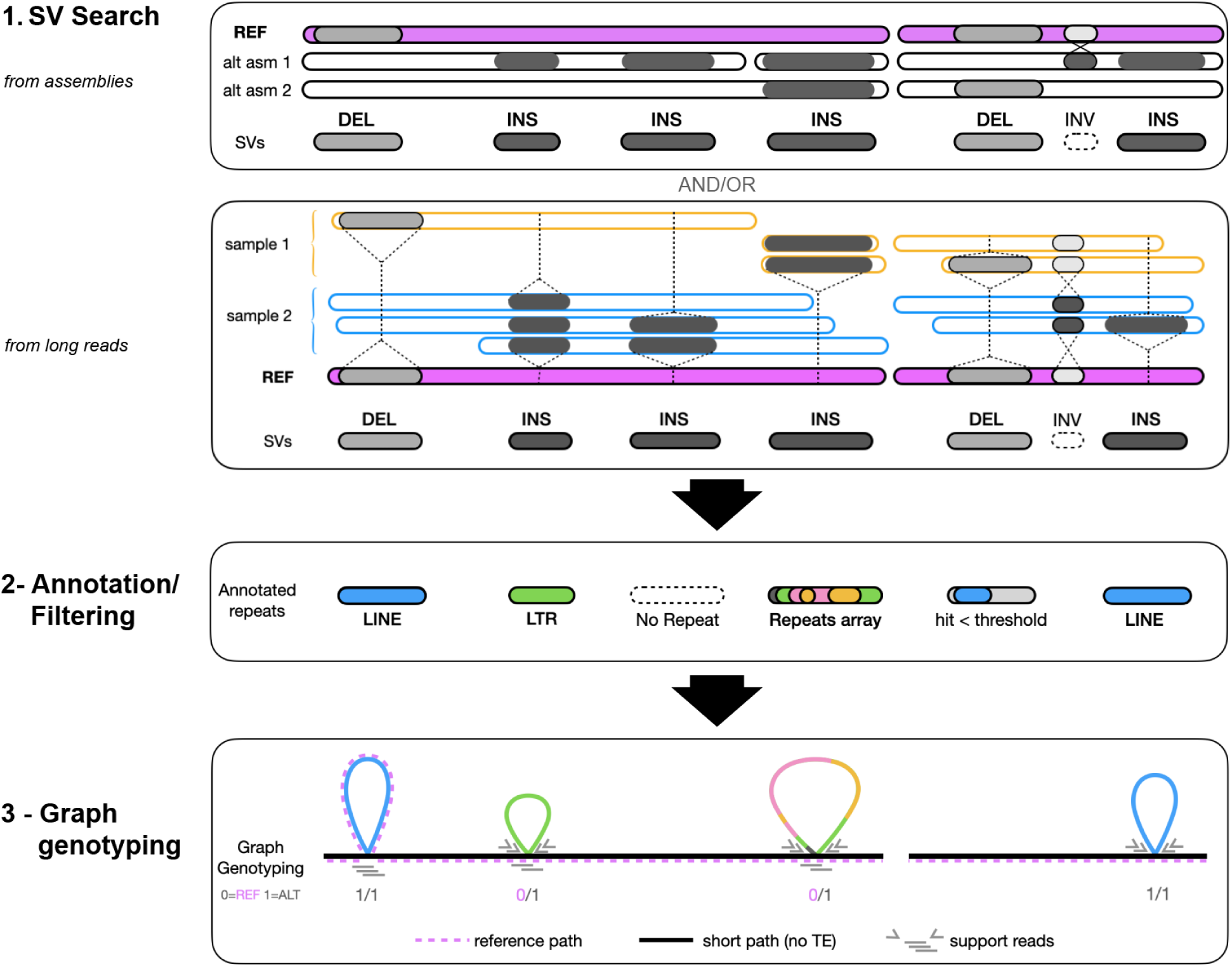
GraffiTE workflow. The pipeline is divided in three main steps: **1- SV search.** Alternative assemblies (“alt asm”, upper box) and/or long-read sets (sample 1, sample 2, lower box) provided in input are aligned to a reference genome (represented in pink) with minimap2. Subsequently, SVs are searched with Svim-asm (from assemblies) or Sniffles 2 (long-read sets) and reported in a VCF file. Insertions (dark gray) and deletion (white) are then retained while other types of SVs (such as inversions) are discarded when using default settings. **2-Annotation/Filtering.** SVs found in different individual assemblies or read-sets, and representing the same loci are merged with the software SURVIVOR. These are further filtered to retain variants which cumulate RepeatMasker’s hit over >80% of their length, based on a user-provided repeat library. **3-Graph genotyping.** Based on annotation thresholds, likely pMEs are retained and a graph genome representing each as a bubble is generated (a.k.a.: TE-graph genome). The reference genome path in the graph (pink dashed line) can either skip (insertions) or include (deletions) a given pME. Finally, short-or long-read sets can be used to genotype each pME using graph aligners bundled in GraffiTE (see Table 1). Additional details are provided in Supplementary Figure 1.

**Table 1.** Software and methods implemented in GraffiTE. GraffiTE steps are labeled according to Figure 1.

| software | version | purpose | GraffiTE step | reference |
| --- | --- | --- | --- | --- |
| minimap2 | 2.24-r1155-dirty | assembly or long-read mapping over a reference genome | 1 - SV search | 50 |
| SVIM-asm | 1.0.3 | SV calling from assembly-to-genome alignments | 1 - SV search | 51 |
| Sniffles2 | 2.0.7 | SV calling from long reads-to-genome alignments | 1 - SV search | 52 |
| samtools | 1.16.1-49-geb703e0 | alignment file sorting | 1 - SV search | 53 |
| SURVIVOR | 1.0.7 | SV merging from individual calls | 1 - SV search | 54 |
| RepeatMasker | 4.1.4 | TE annotation | 2 - Annotation / Filtering | 55 |
| OneCodeToFindThemAll | 1 (adapted) | TE annotation parsing | 2 - Annotation / Filtering | 56 |
| blastn | 2.9.0+ | TSD search module | 2 - Annotation / Filtering | 57 |
| bedtools | 2.27.1 | TSD search module, TE annotation parsing | 2 - Annotation / Filtering | 58 |
| Pangenie | 2.1.0 | Haplotype imputation using k-mers from short-read against the TE pangenome graph | 3 - Graph genotyping | 59 |
| vg | 1.45.0 "Alpicella" | graph induction and genotype calling for Giraffe and GraphAligner | 3 - Graph genotyping | 60 |
| Giraffe | (part of vg) | short-read mapping to TE pangenome graph | 3 - Graph genotyping | 61 |
| GraphAligner | bioconda 1.0.13 | long-read mapping to TE pangenome graph | 3 - Graph genotyping | 62 |
| bcftools | 1.16-81-g8f54545 | vcf file handling and annotation | other | 63 |
| tabix | 1.16-40-g114f5eb | vcf file handling and annotation | other | 64 |

The ability to handle and combine heterogeneous data types (genome assemblies, long- and short-read sets) is a unique aspect of the workflow, and addresses a common scenario of many research projects. For instance, one may use a limited number of genome assemblies or long read sets to establish a high-quality catalog of pMEs, while genotyping them later at population scale using short-reads. In large pangenome projects, assembled genomes are often more convenient to use than raw read sets due to storage limitations. GraffiTE can be directly applied to hundreds of assemblies, and pME detection can be performed without additional data. Another use of GraffiTE can be to annotate and retain pME from a pre-established collection of SVs (stored in a VCF file, see Methods), with the possibility to further perform genotyping on it. Finally, a complete analysis, including pME detection and genotyping can be performed using long-reads set only.

The modular aspect of Nextflow makes it easy to swap or bypass components of the pipeline, and thus will allow seamless upgrade and/or modification with new softwares and routines as needed (Table 1 and Supplementary Figure 1). In addition to their overall performance^49^, the implemented methods were chosen based on several criteria including their availability, being reference-based and using the VCF format to facilitate comparison. We remain attentive to the release of competitive and complementary methods and aim to regularly add new tools and options to the GraffiTE pipeline. Runtime and resources required varies widely according to the type data analyzed, model and computing infrastructure. In order to guide users, we report runtime and resource usage for a variety of analyses and data types, broken down per process (Supplementary Table 4). The program and a detailed documentation (including installation, usage and output description) is available online at https://github.com/cgroza/GraffiTE.

### Simulations demonstrate effective pME detection from different data sources

The first step in the GraffiTE pipeline performs SV search between (i) alternative assemblies and a reference genome, (ii) long-read sets and a reference genome or (iii) both (Figure 1, “1-SV search”). This SV search is followed by annotation of the variants’ sequences and filtering to retain likely pMEs. These analyses produce a VCF file, unified across all samples, listing putative pME presence/absence among samples and their annotations (reported by the “support vector” [SUPP_VEC=] flag of the VCF’s INFO field). To test the ability of the algorithms implemented in GraffiTE to accomplish these tasks, we simulated insertions and deletions of known active families (Alu, LINE-1 and SVA elements) in the chromosome 22 of the human genome and insertions of TIR and LTR elements in the maize chromosome 10 (Figure 2 a, see Methods and Supplementary Methods 1.a.). We introduced background noise in the simulated pMEs by generating random non-pME SVs (insertions or deletions) based on the length distribution of those observed in SV surveys carried out in these species^65,66^. For the human model, all the combinations of tools and data type tested (Figure 2 b, left) yielded both recall (proportion of true pMEs recovered by GraffiTE over the complete set of pMEs simulated) and precision (proportion of true pMEs recovered by GraffiTE over the complete set of calls made by GraffiTE) over 80%. The highest performances were recorded with modes using the assemblies. This is expected since in this experiment the assemblies are simulated 100% complete. In the maize model, long read-only based modes (GT-sn-*) had the lowest score, with recall in the 70-80% range and precisions from 58 to 82%. Meanwhile, all modes using assemblies (GT-*sv-*) had high recall (>94%) and precision ranging between 51 and 84%. Maize is more challenging for GraffiTE due to the fact that ∼80% of the genome is covered by TE sequences, and bears a high level of structural variation^66,67^. In both models, we observed that the lowest scores were recorded when the SV search was performed with Sniffles 2 on simulated PacBIO HiFi reads (GT-sn-HIFI). This was initially unexpected since the HiFi reads are usually more accurate than the ONT reads, but is probably due to the fact that the simulated HiFi reads were shorter on average than the simulated ONT reads, which is also true of real datasets. Thus, while read length was taken into account when controlling for coverage, the read type (ONT vs PacBio HiFi) influenced recall by approximately 5 to 10% in human and maize respectively. Generally, we observed that the precision of the raw outputs can be increased using simple filters: the first filter tested retains only variants with a single hit to a known TE consensus, which increases precision above 80% in the human model and above 58 to 70% for maize (Figure 2 b). The second filter adds a minimum length of 250 bp in humans and 500 bp in maize for a pME to be retained, and allows all combinations of tools and data type to reach their highest precisions, exceeding 90% in humans and reaching between 79 to 86% in maize. The first filter (singe TE hit per variant) had only a limited impact on recall, reduced by <10% in humans and <5% in maize; meanwhile, the second filter (SV length) improved precision without affecting the recall further. The results indicate that model-specific knowledge about the expected mobile TE families is key to improve precision in regards to the detection of true pMEs *vs.* other SV carrying TEs.

**Figure 2.**
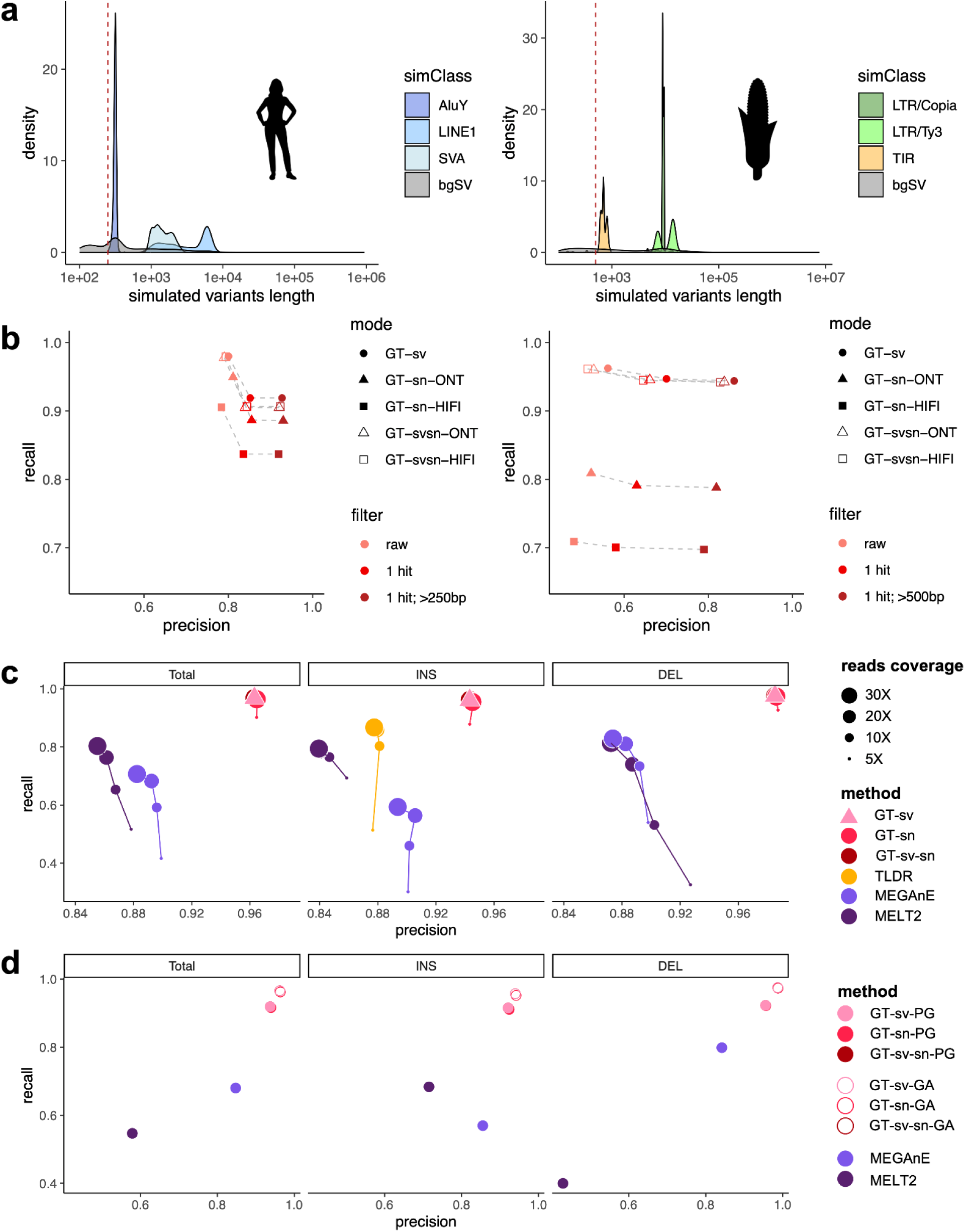
Evaluation of GraffiTE. **a.** Length distribution of the pMEs and background SVs (bgSV) simulated in human (left) and maize (right) datasets. The red dashed line indicates the length cutoff used when filtering variants. **b.** Recall and specificity for pMEs detection on simulated human and maize datasets. “raw”: unfiltered output VCF from GraffiTE, “1 hit”: pMEs annotated with a single RepeatMasker hit representing >= 80% of the variant’s sequence. “1 hit;>250/500bp”: identical to “1 hit” with pMEs < 250 or < 500 bp filtered out. GraffiTE modes: “GT-sv”: GraffiTE-Svim (SV search from assemblies), “GT-sn”: GraffiTE-Sniffles 2 (from long-read sets), “GT-svsn” = GraffiTE Svim-Sniffles 2 (pME search from assemblies and long-read sets, see also Table S1); -ONT suffix: Oxford Nanopore reads; -HIFI suffix: PacBio HiFi reads. See also Supplementary Table 1-B for an extended legend **c.** Evaluation of pME detection performance and comparison with existing tools using a benchmark set derived from the GIAB analysis of HG002/NA24385. Methods legend and acronyms according to Table S1. GT-sv (pink triangle) has no coverage information for pME search, since the variants were directly detected from genomic assemblies of the maternal and paternal chromosomes. Connecting dots indicate runs with increased read coverage. Note that the software TLDR only reports non-reference (INS) pMEs. **d.** Evaluation of pME genotyping performances and comparison with short-read linear-mapping methods using 30X reads coverage. The suffix -PG denotes runs genotyped with Pangenie (short Illumina reads), and -GA GraphAligner (long, PacBio HiFi reads). Methods legend and acronyms according to Supplementary Table 1.

### GraffiTE maintains high performance when benchmarked against real data

The main limitation of the simulations is that the alternative assemblies (simulated chromosomes) are identical to the ground truth, which is unrealistic. Thus, we further tested the relative performance of the tools implemented in GraffiTE for pME detection using a benchmark set derived from^65^ (see Methods). The set includes high-quality pME variants for Alu, LINE1 and SVA elements reported for HG002. Additionally, we applied TLDR^28^, a method designed to find non-reference pME from long-reads, using the long-read sets generated for HG002, and further applied MELT2^6^ and MEGAnE^68^ to short Illumina read-sets. The best performances were obtained with GraffiTE with assemblies (GT-sv), followed closely by GraffiTE modes using long-read sets either alone (GT-sn) or a combination of long-read sets and the diploid assembly of HG002 (GT-sv-sn, Figure 2 c). Compared to TLDR, which searches for pME insertions in long-read data, detection from long reads yielded better performance with GraffiTE, which suggests that Sniffles 2, which performs SV calling in our framework, provides significant improvement for both recall and specificity. Non-surprisingly, short-reads methods for pME detection had the lowest performances, both in terms of recall and precision, as expected when SVs are searched from short-read alignments.

Beyond the detection of pMEs, GraffiTE supports graph-genotyping, i.e. inferring the allele composition for each pME previously detected by mapping short or long reads against a graph genome. We thus tested the performances of Pangenie^59^ (suited for short-reads) and GraphAligner/vg call^62,69^(long-reads) as implemented within GraffiTE. Using the same benchmark set previously, we focus here on the correspondence between the bi-alleleic genotypes provided by the caller and the GIAB genotypes (i.e. the allelic combination at each pME, encoded either 0/0, 0/1 or 1/1 provided for each variants by the GIAB consortium), thus separating heterozygous from homozygous genotypes during testing. We performed graph-genotyping using short or long reads, using the TE-graph genomes created with either GT-sv (assemblies only), GT-sn (long-read only), or GT-svsn (both) (Figure 3 b). We set a fixed read set coverage of 30X for all the tests, as competing methods based on short-reads require to perform both pME detection and genotyping with the same data (MELT2, MEGaNE). The best performances, with precision and recall > 95% were obtained using long-reads and GraphAligner with GraffiTE. As expected, using short-reads for genotyping with GraffiTE (Pangenie) had slightly lower performances, with however, both recall and precision over 90%, outcompeting MEGAnE and MELT2 (note that these two last methods perform pME detection from short reads, unlike GraffiTE, which is expected to reduce the overall recall). Similarly to pME detection, genotyping based on GT-sv (detection from assemblies only) had similar performance than GT-sn (detection from long read) modes.

**Figure 3.**
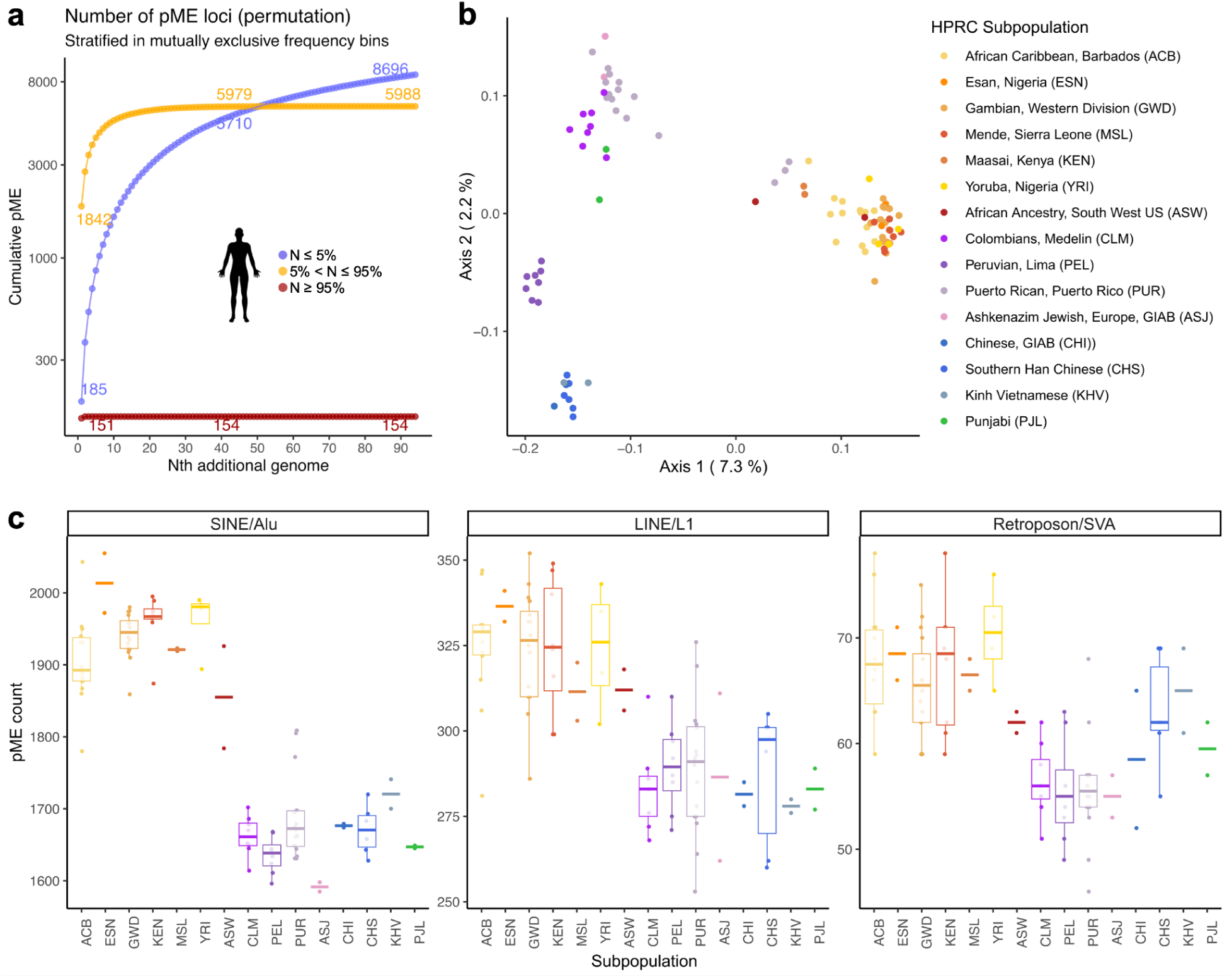
HPRC pME pangenome analysis. **a.** Discovery curves for pMEs detected with GraffiTE using 47 diploid genome assemblies (GT-sv mode) against the reference genome hg38. To reduce false-positives, only pMEs with a single hit on Alu, LINE-1 or SVA and with a size >= 250bp were kept, based on the simulations and benchmark results. For each mutually exclusive frequency bin, the average number of pMEs in N genomes is reported after performing 100 permutations and sampling of the genomes. **b.** PCoA representing the 94 haploid assemblies of the HPRC pangenome using segregating Alu, LINE-1 and SVA insertion polymorphisms. **c.** Count of pMEs for each haploid assembly (N = 94), broken-down per TE type (Alu, LINE-1, SVA) and subpopulation. Bottom and top hinges of the boxes indicate the first and third quartiles of the distribution; the intermediate and thicker bar represents the median. Whiskers extend from the hinges to the largest or lowest value, but no further than 1.5 times the inter-quartile value. Subpopulation codes are referenced according to panel b.

### GraffiTE provides a comprehensive picture of pMEs segregating in the human pangenome

As pangenomes are expected to grow larger in sample representation, the storage and handling of hundreds of raw read sets can represent a computational and financial challenge. With the improvement of both sequencing and assembly methods, it is becoming realistic to perform population-wide study of genetic variation only using assemblies. To illustrate this first use case, we applied GraffiTE to the collection of 47 diploid assemblies recently published by the Human Pangenome Reference Consortium^34^ (HPRC), using the hg38 reference. After the first two steps of GraffiTE (1-SV search, 2-TE filtering), and among 94 haplotypes, a total of 14,838 distinct pME loci were identified, including 11,950 Alu; 2,413 LINE-1 and 475 SVA loci. GraffiTE also reported that 18.7% (451/2413) LINE-1 pMEs bear a signature of 5’ inversion, compatible with twin priming or similar mechanisms^70^. Per individual (diploid genome), the average number of pMEs identified was 3199±322s.d. (2654±274s.d. Alu, 455±43s.d. L1 [incl. 74±7s.d. 5’ inverted] and 90±10s.d. SVA). These estimates are higher than previously reported (2500 to 3000 pMEs per diploid genome) in a recent survey using 30X short-reads datasets from 3,739 individuals from the 1000 Genome Project and BioBank Japan^68^. Additionally, we detected a total of 3100 VNTR polymorphisms located within fixed SVA elements. Using Alu, LINE-1, and SVA pMEs, we show that between 20 and 30 haplotypes are sufficient to recapitulate common segregating variants (haploid genomes frequency > 5%) in the HPRC pangenome (Figure 4 A). Principal Coordinate Analysis (PCoA) using presence/absence call per haplotype recapitulates the characteristic genetic structure of the human population (Figure 4 B). Furthermore, the count of pMEs per subpopulation meets the expectation of higher genetic diversity harbored by individuals of African ancestry (Figure 4 C). In order to highlight the performance of our reference-based approach (minimap2/svim-asm) compared to the method used in^71^ (Minigraph/Cactus), we converted the available graph genome (GFA) to VCF as input for GraffiTE (see Methods). We found that the Minigraph/Cactus approach identifies 788 unique locus in the HPRC pangenome, not detected by GraffiTE however, our approach based on minimap2/svim-asm yielded 6270 unique locus for a union set of 8471 (Supplementary Figure 4).

**Figure 4.**
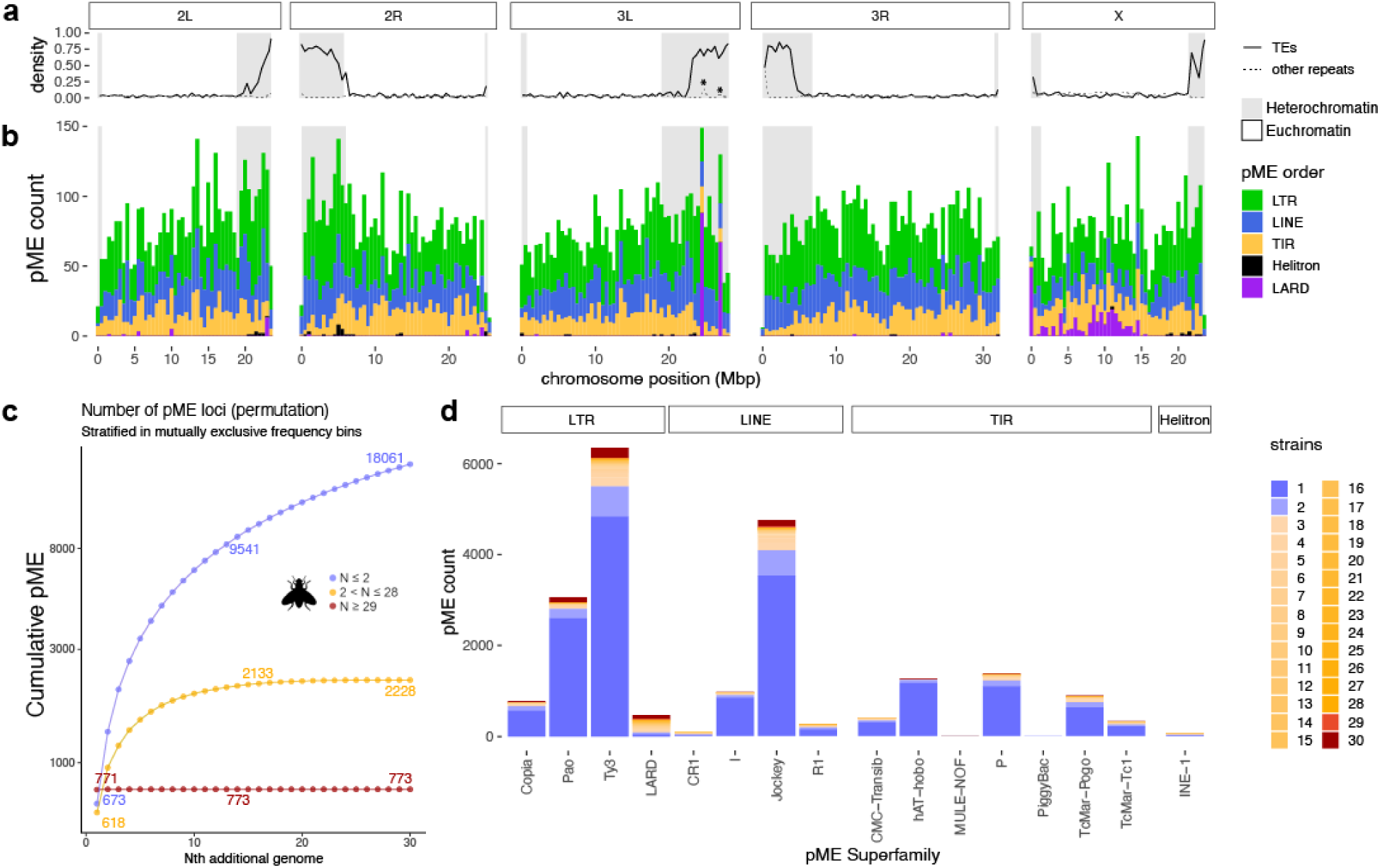
*Drosophila melanogaster* pME pangenome analysis. **a.** Genomic repeat density for TEs and simple repeats (regrouping RepeatMasker annotations: “low complexity”, “simple repeats”, and “Satellite”) as annotated on the reference genome dm6, per bin of 0.5 Mbp. Grey rectangles indicate heterochromatic regions according to Rech et al., 2022. Asterisks represent local peaks discussed in the main text. **b.** Distribution of pME detected by GraffiTE using 30 paired genome assemblies and long-read (ONT) sets, further genotyped with the same long-reads and GraphAligner (GT-sv-sn-GA mode). To reduce putative false-positives, pMEs with more than 1 hit on the TE library were filtered out. pME position is relative to the reference genome dm6; pME counts, per TE order and bins of 0.5 Mbp. Grey rectangle indicates heterochromatic regions according to Rech et al., 2022 **c.** Discovery curves for pMEs detected with GraffiTE. For each mutually exclusive frequency bin (blue: shared by 1-2 genomes, yellow: 3-28, red: 29-30), the average number of pMEs in N genomes is reported after performing 100 permutations and sampling of the genomes. **d.** Count of pMEs detected by GraffiTE according to their superfamily and colored by their frequency in the pangenome.

### Analysis of the *Drosophila melanogaster* pangenome confirms a highly dynamic repeatome

*D. melanogaster* displays a vastly different context for pME detection compared to humans, as the overall genomic TE content is rather small (∼20% of the genome) but a much greater number of TE families are mobile, and many do so in a population specific manner^11^. We applied GraffiTE to a subset of 30 *Drosophila melanogaster* genomes produced and analyzed by Rech et al.^8^. These genomes encompass an identical number of strains originating from 12 distinct geographic locations, chosen based on their shared utilization of ONT long-read sequences. Genome assemblies and raw, long-read sets, were used as input for GraffiTE (GT-svsn mode) and graph genotyping was performed with Graphaligner/vg call on the long-read (GT-svsn-GA mode). This combination of tools showcases an analysis taking advantage of all the data available, and in addition to the analysis of the human pangenome, provides bi-allelic genotype for each pME in each strain represented in the pangenome. The output VCF was further filtered to retain variants with a single RepeatMasker hit on the authors’ manually curated TE library (MCTE), as our simulations show that this simple filtering improves the method’s specificity (Figure 2 a). Furthermore, we excluded variants for which the GraphAligner/vg call genotypes were fixed for all samples relative to dm6, and those with missing genotypes. Because fundamental differences in the reporting of events between GraffiTE and ^8^ prevented us to perform an unbiased comparison of the pME found by each method (Supplementary Methods 2 b and Supplementary Figures 5-11), we rather focused on expanding the pME search and genotyping to heterochromatic regions, which were not originally analyzed. After graph-genotyping using ONT long reads, GraffiTE reported a total of 21,062 pMEs, including 15,581 euchromatic and 5,481 heterochromatic variants (Figure 4, a-b). Genotyping of the same TE-pangenome with available short-reads and Pangenie (GT-svsn-PG mode) only yielded 10,539 variants (using the same filters as above), including 9,464 reported with long-reads (Supplementary Figure 12). Discovery curves (Figure 4 C) indicate that approximately 20 genomes are sufficient to discover the most common polymorphisms (pME loci detected in 2 < N < 29 genomes relative to the reference ISO-1/dm6), matching the previous report in euchromatic regions^8^. This comparison suggests that the discovery rate of GraffiTE in heterochromatic regions might be comparable to the euchromatin in this *Drosophila* pangenome. Nevertheless, the bulk of pME appears to be segregating a very low frequency, 18,061 variants being found in 1 to 2 genomes only (Figure 4 c,d). This result is largely expected given the documented TE variation observed in natural *D. melanogaster* populations^11,72^. In contrast to other pME families, we noticed an uneven distribution of variants for members of the INE-1 (Rolling-circle/Helitron) and LARD elements. Most novel INE-1 variants are found in heterochromatic regions, so does LARD insertions, which are also found vastly through the euchromatin of the chromosome X. Based on rather high frequencies on the pangenome (Figure 4 d), and the fact that these families are estimated to be much older than the other pMEs^8^, it is likely that these variants are indicative of the lack of completion of the dm6 reference in traditionally hard to access regions. In addition, we notice that LARDs distribution shows two unusual peaks in the heterochromatic regions of chromosome 3L. Both peaks coincide with elevated density of non-TE repeats (low-complexity, simple repeats and satellites, highlighted with asterisks, Figure 4 a). These may be caused by two non-exclusive factors: (i) the lack of completion of the dm6 reference in these regions, and (ii) possible false positive hot-spots, as repeat-rich regions remain challenging for genome to genome and long-read to genome SV search methods implemented in GraffiTE^73^.

### Exploration of an edge case: extreme variation at the *bz* locus in Zea mays

To test the performances of GraffiTE in challenging models, we analyzed the genomic variation at the *bz* locus in a 23-samples pangenome of *Zea mays*. We first ran GraffiTE in GT-sn mode (long-read samples *vs.* reference), using 23 read sets available for the same number of varieties, sequenced by^74^ (three read sets were not used as they did cause errors with the onboard tools). Reads were aligned against the recently published telomere-to-telomere genome of the strain Mo17 (Zm-Mo17-T2T)^75^. We favored this reference as the backbone of our analysis due to its high contiguity and completion. We used the VCF produced by GraffiTE to generate a GFA (graph genome) file for the locus *bz* (Zm-Mo17-T2T CM039158.1:13076986-13149277), as well as the TE annotation produced by our pipeline (see method) as an input for Bandage^76^, in order to produce a graphical layout. We identified 13 variants (Figure 5 a, variants 1-13), whose frequencies in the pangenome range between 1 (variant 4: Sniffles2.INS.1940M8 [LTR/Copia], variant 7: Sniffles2.INS.1941M8 [LTR/Copia] and variant 10: Sniffles2.INS.194AM8 [multiple TEs]) and 15 (variant 9: Sniffles2.INS.1949M8 [TIR/DTH], Figure 5 a, Table 2). Upon manual inspection (blastn dot-plot) we noticed that all the variants that include copies of LTR/Copia (variants 1,4,7,8,10 and 12) were full-length LTR elements (Supplementary Table 5), likely the product of recent insertion polymorphism (i.e., pMEs). None of the other variants showed structural features from their classes which would be compatible with *bona fide* pMEs, i.e.: no TIR nor LTR could be found at their boundaries, suggesting that these are non-pMEs SVs carrying pieces of older elements. A known polymorphism for *Grande1*, an LTR/Ty3 retrotransposon^77^, is not present in this pangenome, however two nested structural variants representing internal deletions of this element are observed (variants 5 and 6). Though GraffiTE captured the TE sequences correctly, the annotations given from the pan-EDTA library were in some instances fragmented and/or attributed to 2 or more different consensus sequences (Table 2 and Supplementary Table 5), highlighting the importance of manual curation and model-specific knowledge during the interpretation of GraffiTE results. In order to further compare GraffiTE results to reference datasets, we used the blastn functionality of Bandage (after merging all possible nodes and using a seed of 22 nucleotides, see Methods) to map 8 *bz* haplotypes described in detail by^77^. Polymorphisms at the 13 SVs (including the 6 LTR/Copia pMEs) are clearly apparent (Figure 5 b), as well as additional polymorphisms not captured in the 23-samples pangenome (i.e. not represented by a bubble). For example, the polymorphisms of “*Grande1*”, a LTR/Ty3 retroelement (delineated by triangles in Figure 5), are in complete agreement with the description of^77^. p

**Figure 5.**
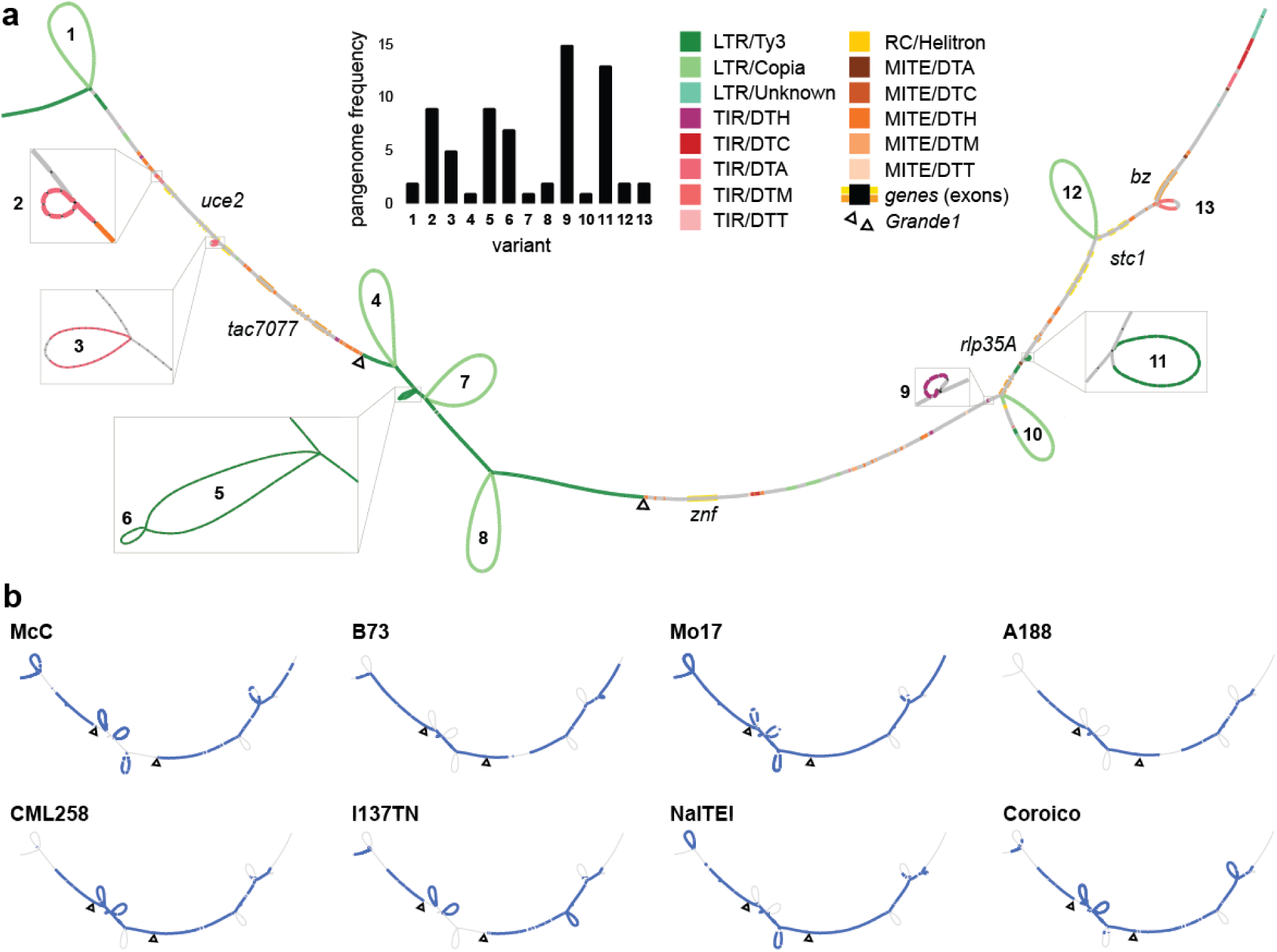
Structural variation at the *bz* locus in *Zea mays*. **a.** Bandage representation of the *bz* locus using the VCF and annotations provided by GraffiTE. The graph resolution (nodes size) is 32 bp. Bubbles in the graph represent each structural variant or pME detected (see table 2). The histogram reports each variant frequency in the pangenome. Shared paths (non-bubbles segments) were annotated using RepeatMasker. Genes’ exons are represented by larger yellow or orange highlights. The region of the graph for the *Grande1* LTR/Ty3 retrotransposon is indicated between the triangle markers. **b.** Mapping of the 8 *bz* haplotypes described by Dooner 2006 over the 23-samples pangenome with Bandage. Colored segments represent DNA regions present in each haplotype. The polymorphism at *Grande1* is shown with triangle markers. Unconnected blue bubbles (i.e. not connected to a shared path of the graph) are attributed to spurious blastn hits, originating from homologous TE present in the query haplotype, but absent from the 23-samples graph.

**Table 2.** Description of the 13 SVs and pMEs discovered by GraffiTE in a 23-samples *Zea mays* pangenome. INS: insertions relative to Zm-Mo17-T2T; DEL: deletion relative to Zm-Mo17-T2T; hits: number of distinct RepeatMasker hits within the variant’s sequence; frag.: number of fragments per hit. strand: strand of the RepeatMasker hit(s) relative to the consensus sequence(s).

| Structural Variation |  |  |  |  | TE annotation |  |  |  |  |  |
| --- | --- | --- | --- | --- | --- | --- | --- | --- | --- | --- |
| # | GraffiTE ID | type | length (bp) | samples with variant (23-pangenome) | hits | frag. | consensus name | class | strand | Manual annotation |
| 1 | Sniffles2.INS.1919M8 | INS | 9127 | Ms71,Oh43 | 2 | 1,1 | chr9_D_32444039<br>ji_AC200613_6936 | LTR/Copia | +,+ | LTR/Copia pME |
| 2 | Sniffles2.DEL.1930M8 | DEL | 204 | CML52,CML69,CML103,CML322,CML333,II14H,Ky21,M162W,NC358 | 1 | 1 | DTA_ZM00052_consensus | TIR/DTA | + | SV containing TE fragments |
| 3 | Sniffles2.DEL.1931M8 | DEL | 956 | CML69,CML322,HP301,Ky21,M162W | 1 | 2 | DTA_ZM00148_consensus | TIR/DTA | - | SV containing TE fragments |
| 4 | Sniffles2.INS.1940M8 | INS | 8947 | M162W | 1 | 1 | opie_AC198924_6206 | LTR/Copia | + | LTR/Copia pME |
| 5 | Sniffles2.DEL.195BM8 | DEL | 1809 | HP301,II14H,Ki3,Ky21,Ms71,NC358,Oh43,Oh7B,P39 | 1 | 3 | grande_AC200214_6803 | LTR/Ty3 | - | SV containing TE fragments |
| 6 | Sniffles2.DEL.195CM8 | DEL | 293 | CML69,CML103,II14H,Ms71,Oh7B,P39,Tx303 | 1 | 1 | grande_AC200214_6803 | LTR/Ty3 | - | SV containing TE fragments |
| 7 | Sniffles2.INS.1941M8 | INS | 8843 | CML103 | 2 | 1,1 | chr4_D_40959532<br>chr8_P_116327109 | LTR/Copia | +,+ | LTR/Copia pME |
| 8 | Sniffles2.INS.1942M8 | INS | 9410 | CML228,Ki11 | 2 | 1,1 | ji_AC215728_13156<br>ji_AC200613_6936 | LTR/Copia | +,+ | LTR/Copia pME |
| 9 | Sniffles2.INS.1949M8 | INS | 183 | CML52,CML103,CML228,CML247,CML277,CML322,HP301,II14H,Ki11,M162W,Mo18W,NC350,Oh43,Oh7B,Tx303 | 1 | 1 | DTH_ZM00021_consensus | TIR/DTH | - | SV containing TE fragments |
| 10 | Sniffles2.INS.194AM8 | INS | 8033 | Ms71 | 6 | 3,1,1,1,1,1 | debeh_AC177840_246,<br>TE_00010160<br>TE_00004946<br>TE_00010537_INT<br>name_AC197689_5725<br>TE_00009602_INT | LTR/Copia<br>TIR/Helitron<br>TIR/DTM<br>LTR/Ty3<br>LTR/unknown<br>LTR/Copia | -,+,+,+<br>+,- | LTR/Copia pME |
| 11 | Sniffles2.DEL.1962M8 | DEL | 686 | CML69,CML247,CML277,HP301,II14H,Ki3,M37W,Mo18W,Ms71,NC358,Oh43,Oh7B,P39 | 1 | 1 | cinful_zeon_AC198933_6214 | LTR/Ty3 | - | SV containing TE fragments |
| 12 | Sniffles2.INS.194EM8 | INS | 8847 | CML69,M162W | 1 | 1 | ji_AC204382_8228 | LTR/Copia | + | LTR/Copia pME |
| 13 | Sniffles2.INS.1951M8 | INS | 2558 | CML247,CML277 | 1 | 3 | Zm01468_AC186160_1 | TIR/DTM | + | SV containing TE fragments |

### GraffiTE detects segregating repeat families in a model with limited prior knowledge about TEs

We previously demonstrated two use cases of GraffiTE in model species for which extensive knowledge about their TE biology is available. In this final use case, we demonstrate that GraffiTE can also be applied to new models, with little-to-no information available about their TEs. *Cannabis sativa* is an emerging crop of economic and medical importance, and for which genomic resources are accumulating, including several pangenome projects^78^. Moreover, *C.sativa* is an example of TE-rich genomes (>70% of the genome^79^) for which high-level of polymorphism is expected^80^. Nine publicly available assemblies were used as input for GraffiTE (GT-sv mode) using Cs10 as reference. In contrast to human and *Drosophila* models, which use manually curated libraries, putative pME annotations were often composed of multiple hits against different consensus sequences from the automatically generated TE library. This is expected for consensus sequences lacking manual curation of TE catalogs. To circumvent this limitation, we further extracted and clustered together the sequences of all 75,679 variants present in the raw TE-graph genome generated by GraffiTE (Figure 6 a and Methods). We then retained clusters with at least 3 distinct loci in the TE-graph genome, and with a size between 200 and 40,000 bp as probable pMEs and computed the discovery curves (Figure 6 B). We deliberately applied a loose filter on variant length in order to be able to detect the largest type of TE known, such as Maverick/Polinton^81^; it is nevertheless important to note that this may yield false positives, such as repeated, chimerical variant sequences containing TEs. We showed that 7 assemblies were required to observe the common pMEs in the *C. sativa* pangenome (pMEs found in 2 to 8 assemblies), while 208 variants from the reference Cs10 were shared by all 9 alternative genomes. Both common and rare (singleton) pMEs were extremely abundant, for a total of 42,517 loci (in contrast, ∼15,000 and ∼20,000 pMEs were found in 94 human and 30 *D. melanogaster* assemblies, respectively). Adding long read sets available for 5 of the assemblies to the original analysis (GT-svsn mode) further increased the set of candidate pME by 19,207 loci (Supplementary Figure 13). Thus, for comprehensive discovery, we encourage users to use as much of both long-reads and assembly available. To further illustrate how GraffiTE can be used to improve TE annotation, we selected the most abundant pME clusters (putative TE families) and measured their abundance in the pangenome (Figure 6 C). Based on copy number, the 100 most abundant pME families represent approximately 25 Mbp of facultative DNA sequence, differentially present or absent between the 9 assemblies and the reference Cs10; the most interspersed TE family (representative sequence: Pink_pepper.svim_asm.INS.10081) being represented by a total of 756 loci in the pangenome. Manual annotation for a subset of these families highlight the diversity of the (recently) active TEs in *C. sativa*, including Miniature Inverted-repeat Transposable Elements (Class II, MITE), LTR Retrotransposons (Class I) and DNA/TIR of the MULE-MuDR superfamily (Class II).

**Figure 6.**
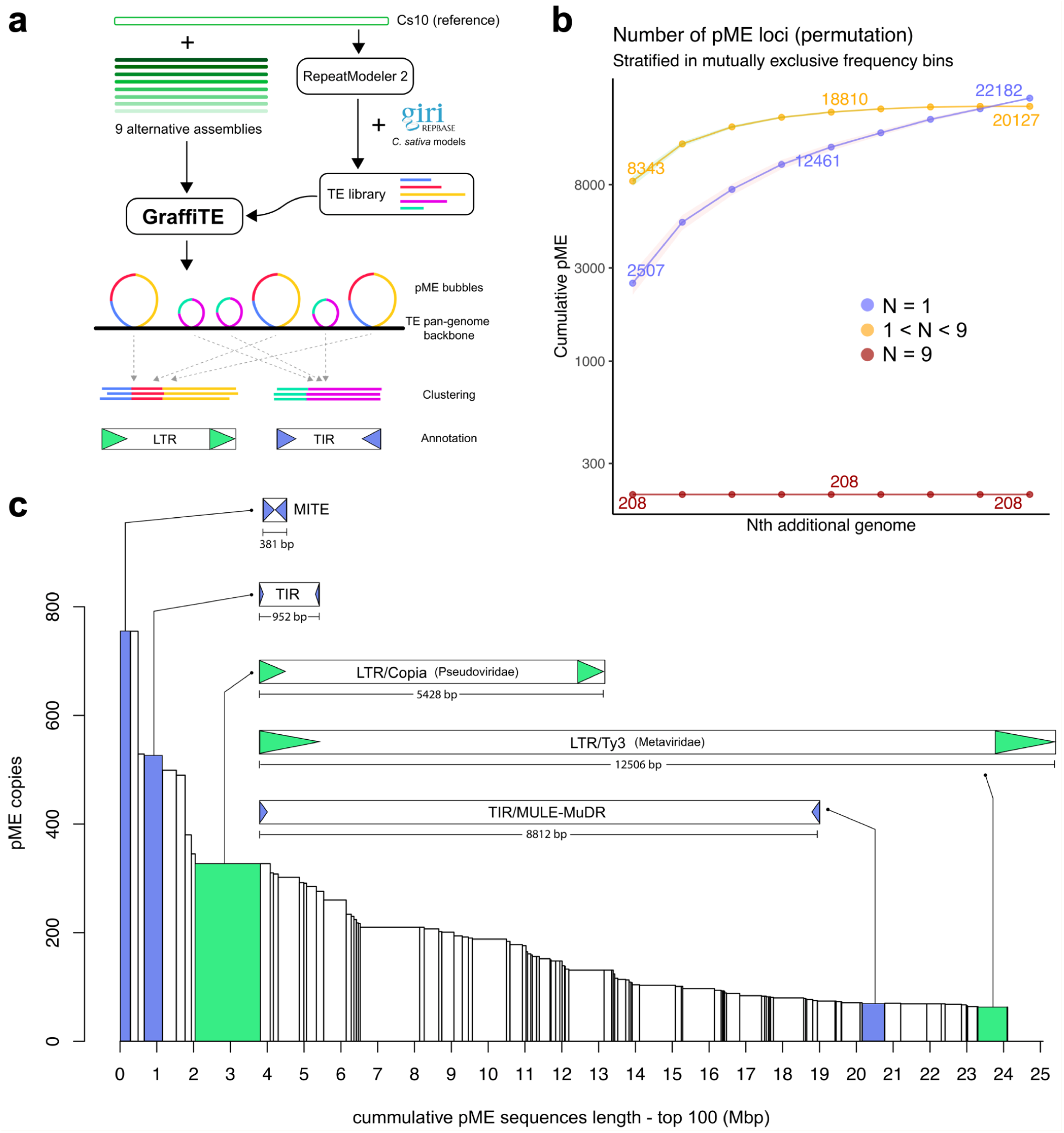
*Cannabis sativa* pME pangenome analysis. **a.** Overview of the experiment: the reference genome Cs10 was used to generate an automatic TE library with RepeatMolder 2 and subsequently merged with a collection of manually curated consensus sequences from Repbase. Next, 9 alternative assemblies were used as input for GraffiTE (GT-sv mode). Most of the discovered candidate pMEs contained multiple TE annotation from the automatic library; in order to reconstruct TE families, the sequences of 75,679 putative pMEs were clustered with MMseqs2 and variants belonging to clusters with 3 or more sequences were kept for analysis. **b.** Discovery curves for the selected pMEs in 9 genome assemblies, relative to the reference Cs10. **c.** Distribution of pME clusters size, in total number of loci (y axis) and total sequence length in the pangenome (x axis = pME loci number x pME lengths per cluster). The manually annotated sequence of 5 representative pME families are shown to illustrate the diversity of TE segregating between *C. sativa* cultivars.

## Discussion

We created GraffiTE with the goal of facilitating pME analysis in a wide-range of organisms, enabling the exploration of TE dynamics in model species with or without prior information about their mobilome. A particular emphasis was made to include a collection of methods ready for the pangenome era, allowing the direct detection of pMEs from multiple genome assemblies and/or long-read sets. More importantly, GraffiTE introduces for the first time the ability to perform graph-based genotyping of pME loci, by leveraging cutting-edge software such as Pangenie^59^ and GraphAligner^62^.

A major goal was to provide flexibility and ease of use for a wide audience of researchers. To do so, GraffiTE relies on Nextflow, which provides multiple avantages when designing such pipelines including: (i) the self-containment of the code and its dependencies using a single Apptainer (formerly known as Singularity) image, (ii) the ability to swap or combine different software and modules, and (iii) the scalability of the pipeline, allowing its deployment to a wide array of systems including high performance clusters (HPC) and cloud services. Furthermore, GraffiTE provides detailed annotated outputs, and is released with extensive documentation.

Another strength of GraffiTE is the wide variety of data types to which it can be applied. pMEs can be detected from genome assemblies or any type of long-read data, and genotyping can be performed using short- and long-read sets. This flexibility allows researchers to get the most out of their data; for example, by performing the initial SV search with high-quality – though perhaps less abundant – data, such as chromosome-level assemblies and long-read sequences, while genotyping in larger cohorts or populations using cost-effective short-read sets. Users should consider the inherent limitations of each type of data. For SV discovery, assemblies may be incomplete and thus their quality should be taken into consideration (though in some cases assembly have been shown to perform better than long-reads^49^). Short-reads genotyping will reduce recall (i.e. some variants detected with long reads or assemblies may not be able to be genotyped with short reads due to their length and structural complexity); this consideration is important when considering rare variants.

GraffiTE provides additional features not found in existing software, such as the ability to report deletions (pMEs present in the reference genome but absent from a given sample), a feature absent from long-read methods such as TELR^29^ or TLDR^28^. While TrEMOLO^82^ recently introduced the ability to detect pMEs from an alternative genome assembly, GraffiTE expands this feature to multiple genomes in a single analysis. Furthermore, GraffiTE uses and produces annotated VCFs as a standard, which allows swift interoperability both between tools within the pipeline but also for the users to share and compare results. Finally, users can annotate and genotype pME from SV call sets generated by other tools, as long as both reference and alternative alleles are documented in an input VCF. Thus, GraffiTE can be directly used on already established collections of SVs, and will allow filtering and annotation of variants to retain and graph-genotype pMEs.

Simulations show that the different modes (pME detection from assemblies only, long-read only, or combined) produce high recall (low false negatives) and that specificity and precision can be greatly improved with simple hypothesis-driven filters, such as pME length and type of annotation. In real data derived from the Genome in a Bottle (GIAB) benchmark^65^, all modes of GraffiTE yield results above the scoring of alternative methods such as TLDR (long-reads) or MELT2 and MEGanE (short-reads), including when using only the paternal and maternal assemblies of HG002. We note that these assemblies are exceptional given their near completion<u>(Rautiainen et al. 2023)</u>, and other assembly completeness may vary and will directly affect the tool’s ability to detect SVs.

We further evaluated the performance of graph-based genotyping, showing a high concordance with the high-quality bi-alleleic genotypes reported by the GIAB consortium and GraffiTE (Figure 2 c). This result demonstrates that graph-genotyping improves genotype quality of pME variants, which is particularly relevant when pMEs are used in association studies such as GWAS or molecular QTLs^19,83^. Applying GraffiTE with genome assemblies only remains highly informative as we show that 20 to 30 human haploid assemblies are sufficient to discover common polymorphisms in the HPRC pangenome^34^, without the need for raw long-read data. This is particularly relevant, as the storage of raw data from larger pangenomes can be a significant limitation for research teams.

Application to a *Drosophila melanogaster* pangenome highlights the flexibility of the pipeline to adapt to various models with very diverse TE dynamics. In particular, our analysis shows great consistency with population genetics reports of TE polymorphisms in this species^8,11,72^. Furthermore, GraffiTE fared well in notoriously challenging systems, such as *Zea mays*. Our analysis of 23 long-read samples from multiple cultivars allowed us to tease apart *bona fide* pMEs (full-length LTR/Copia) from other SVs carrying TE pieces at the *bz* locus. In our last use case, we illustrated with a *C. sativa* pangenome how GraffiTE can be used in non- or emerging model species, for which little-to-no information about TE is available. A simple clustering-based strategy quickly identified the most abundant polymorphic repeats, which could be further manually annotated. Thus, we argue that in complement to automated TE discovery approaches, such as RepeatModeler^84^, REPET^85^ or EDTA^86^, GraffiTE can be a useful addition to library curation tools by identifying the precise boundaries of the most recent (polymorphic) TE families, an approach also recently implemented in Pantera^87^.

As with any automated approach, we can identify current limitations with the proposed software. SVs annotation, in order to retain likely pMEs, remains particularly challenging. Our current implementation based on homology between the variant sequence and a known TE library, favors generalization but can occasionally provide erroneous or ambiguous results. For example, full-length LTR insertions (proviral LTR) may be reported as three separate hits (5’ LTR, internal sequence, 5’ LTR) if the library doesn’t report the relationship between LTR and internal consensus. Though we attempt to mitigate this behavior using OneCodeToFindThemAll^56^, recombination between different LTR families during transposition may lead to similar patterns and cannot be easily detected automatically. In very rare cases, we found that RepeatMasker can report hits much longer than the consensus sequence, which can be incompatible with the adjudicated pME family. Such artifacts can be later discarded by the user using the detailed output of GraffiTE, applying specific knowledge about the concerned TE families. We also noted that with some scaffolded genomes, minimap2 can return an error due to a CIGAR string being too long. This is caused by large stretches of N residues used to patch the scaffolds, and we provide an option to automatically split the input contigs in case this error occurs (see GraffiTE documentation). Finally, the accuracy of genotyping pMEs segregating within other pMEs (variants in variants) has not been tested. Though rare, if such a case presents itself, GraffiTE will output two variants for the same positions; we invite users to look for this possibility by reviewing the provided annotations at loci sharing the same coordinates. Here, we have introduced GraffiTE, a powerful and user-oriented tool that facilitates pME analysis in a wide range of organisms. It enables precise analyses in model species as well as exploration of TE dynamics in systems with limited mobilome information. With its graph-based genotyping capabilities and compatibility with multiple data types, including genome assemblies and long-read datasets, GraffiTE offers flexibility and optimal use of researcher’s ressources. The software provides unique features such as analyzing multiple assemblies at a time, annotation of structural variants, and interoperability through annotated VCFs. It demonstrates high performance and accuracy in TE detection and genotyping, making it valuable for association studies and pangenome analyses. We aim to provide continuous support for the pipeline, through its Github page (https://github.com/cgroza/GraffiTE) and are looking forward to incorporating user’s feedback into updated versions. Future work is planned to tackle complex TE polymorphisms, such as LTR-LTR recombination, pME nesting, and perform gold-standard method evaluations through PCR assays. Thus, we believe that GraffiTE represents a valuable contribution to TE research, enabling insights into TE biology, genome evolution, and molecular diversity.

## Methods

### Implementation

#### SV search

The first step of GraffiTE searches for SV between a reference genome and one or more alternative assemblies for the same species. SV can also be searched from long-read sets (either alternatively or in combination with search from assembled genomes). When using assemblies, each input file is expected to be representing a single haplotype in fasta format (if diploid assemblies are used, each haplotype is analyzed separately). Each queried assembly is first aligned onto the reference with minimap2 (v.2.24-r1155-dirty) using the preset -x asm5 which is optimized for genome-to-genome alignment below 5% divergence (minimap2 documentation, accessible online at: https://lh3.github.io/minimap2/minimap2.html). This default preset can be modified to accommodate more divergent data, up to 20% divergence (see GraffiTE documentation). Additional parameters -a --cs -r2k are also applied, as recommended for the next program SVIM-ASM (v.1.0.3) which calls SVs based on minimap2 outputs (BAM alignments). By default SVIM-ASM is tasked to report insertions and deletions >= 100 bp relative to the reference genome (inversions, duplication and translocation are not reported in order to focus on candidate products of transposition). From long-reads, SVs are searched by first mapping reads to the reference genome with minimap2 (presets -x ont/pb/hifi are used according to the input reads, see online documentation). Then, insertion or deletion variants >= 100bp are searched in the alignments using Sniffles 2 (v.2.0.7). Upon SV calling, variants are reported in one VCF file per alternative assembly or read set. If multiple samples are used, individual VCFs are further merged using SURVIVOR (v.1.0.7). SVs are merged into the same loci if they belong to the same type (INS or DEL), their distance is within 10% of their length and each SV >= 100bp (SURVIVOR merge 0.1 0 1 0 0 100). GraffiTE allows variants found in only one assembly or read-set to be kept (singletons).

This SV search can be bypassed by directly providing a sequence-resolved VCF (REF and ALT sequence for each variant must be provided) to GraffiTE (Supplementary Figure 1).

#### SV filtering

The next step for GraffiTE is to filter the candidate SVs to retain likely pMEs. Insertion and deletion sequences are extracted from the merged VCF (or alternatively a provided sequence-resolved VCF) in FASTA, and scanned using RepeatMasker v. 4.1.4^55^ using its RMBlastn engine (modified Blastn distribution). At this step, a user-provided fasta library of TE models to analyze (consensus sequences) is required. The input TE library can be the full collection consensus sequences for the TEs of the target species, or only include TE families of interest. Since TEs can be hitchhiking within SVs caused by other events than transposition, GraffiTE requires that TEs identified by RepeatMasker covers >= 80% of the variant sequence. Using this threshold, the retained variants can include a single or multiple hits against different known TEs. In some cases, multiple hits within the same SV can be due to rearrangement of the newly inserted TE sequence (insertions, deletions or inversion in the TE copy relative to the consensus sequence), others source of multiple TE hits within SV can be caused by distinction in the TE library between long terminal repeat (LTR) and internal part of LTR elements, as well as artifacts (e.g. spurious hits) caused by the RepeatMasker algorithm. To help identify and regroup hits corresponding to the same TE copy (fragments), GraffiTE uses an adapted version (correcting regular expression searches) OneCodeToFindThemAll.pl^56^, a Perl script dedicated to this task. Note that to take the best advantage of this step, it is preferable that the nomenclature of the TE library follows the RepeatMasker style: TENAME#Order/Superfamily (e.g., L1HS#LINE/L1) and that internal (l) and LTR sequences of the same elements are named such as TENAME_I#LTR/Superfamily and TENAME_LTR#LTR/Superfamily (e.g., ROO_I#LTR/Pao and ROO_LTR#LTR/Pao). Following this step, for each SV retained the number of hits (distinct TE instances), fragments (pieces of the same TE instance) and classification details are reported in a VCF file.

#### TSD search

We implemented a module to search for target site duplications (TSD) which can be the hallmark of transposition in many TE families. Any TE-containing variant retained following the previous step (SV filtering) and for which a single TE is identified (this can include hits made of multiple fragments) will be processed. The TSD (if present) is often located at the 5’ or 3’ end of the SV sequence, while the second TSD is present at the opposite end, in the flanking region. However, TSDs can sometimes be found overlapping the junction between the flanking and the first base pairs of the SV. To take these cases into consideration, the TSD module of GraffiTE first removes the base pairs masked by RepeatMasker in the SV, leaving (if any reminder) a few base pairs at the 5’ and 3’ end. Then, these reminders (if any) are concatenated to 30 bp of flanking sequence (directly adjacent to the SV breakpoints) at each respective end (Supplementary Figure 2 a). These reconstructed flanking sequences are then compared to each other using blastn (v. 2.9.0+), with a seed of 4bp (thus the minimal TSD size reported is 4bp). Though the best match (longest hit with the least mismatches) for each comparison (each pME) is reported in an output table, we implemented an empirical filter to reduce false positives. The filter returns a PASS flag (and report the TSD sequence in the output VCF) if the TSD initiates or ends within 5bp of the defined TE end (according to the consolidated RepeatMasker outputs) and if the number of mismatches is below 1 or [TSD-length]xDIV whichever the larger, with DIV the divergence of the TE copy relative to its consensus, as reported by RepeatMasker (i.e. older TE insertion tolerates more mismatches). An additional subroutine makes sure that the TSDs are located outside any poly-A (or T) tail (Supplementary Figure 2 b).

#### Additional annotation filters

Initial testing on human data revealed that the automated annotation of GraffiTE was, as-is, likely to mislabel specific types of TE-associated polymorphisms. In particular, we noted that 5’ LINE-1 inversions (due to twin-priming and associated mechanisms^70,88^) would be reported as two distinct L1 insertions on opposite orientations and are not stitched together by OneCodeToFindThemAll. Accordingly, we implemented a filter to recognize such configuration and re-label insertions as a single “twin-primed” L1 (Supplementary Figure 3). In addition, GraffiTE is able to detect variation in the number of tandem repeats (VNTR) present within SINE-VNTR-Alu elements (SVA), a human lineage-specific type of TE. In order to distinguish insertion polymorphism of new SVA copies (presence/absence of the SVA) from VNTR-only variation (SVA present in both haplotypes, but with VNTR length variation between alleles) the variant’s sequence (being either an insertion or a deletion relative to the reference genome) is blasted against the consensus sequence of each of the 6 known SVA models (SVA_A, B, C, D, E and F). If the variant fully maps within the VNTR region of the consensus, it will be labeled “VNTR-only” in the INFO field of the output VCF, while canonical SVA insertion (i.e. pME) will not harbor this tag.

#### Variant validation and genotyping

Following the SV detection and annotation to retain likely pMEs, GraffiTE gives the possibility to validate and genotype (i.e. predicts the alleles of) each variant by mapping reads from individual samples onto a genome graph where TE-containing haplotypes (each insertion or deletion sequence in the annotated VCF) are represented as bubbles and TE-absent haplotypes are represented by short paths (skipping the TE bubbles). In order to accommodate different sequencing technology, we included Pangenie (short-reads^59^), Giraffe (short-reads^61^) and GraphAligner (long-reads, including low-fidelity ONT^62^).

### Evaluation of GraffiTE performance

All evaluations were performed using GraffiTE v 0.2.3 (02-21-22, GitHub SHA: 557f6402dbe9669af30081270cbe170fbf044477)

#### Simulations

In order to evaluate the ability of the tools implemented to retrieve pMEs from either assemblies or long-read sets, we simulated pMEs and random (background) SVs on random 1Mb stretches of either the human chromosome 22 or the maize chromosome 10. For each simulation, 20 pMEs were randomly sampled among members of the AluY, L1HS/L1PA2 and SVA_E,F superfamilies in human (as present in the reference genome GRCh38.p14/hg38) or among members of the TIR/DTA, LTR/Ty3 (formerly known as Gypsy) and LTR/Copia superfamilies in maize (as present in the reference genome Zm-Mo17-REFERENCE-CAU-2.0). For each simulated pME a realistic target site duplication (TSD) is also produced (see Supplementary Methods 1). In addition to pMEs, and for each simulation, 20 background SVs were sampled from random intervals in the genome (Supplementary Methods 1a). Our simulator produces VCF files that are then passed to SimuG, a perl script tasked to create artificial genomes based on the variants described in the VCF^89^. Once generated, each simulated genome was artificially sequenced at 10X depth with PBSIM3^90^.

#### GIAB Benchmark

An ideal benchmark dataset would include a set of TE insertion polymorphisms for which the presence or absence is known at the allelic level with high-confidence (ideally PCR-based genotyping) in multiple individuals. Such an optimal dataset does not directly exist for pMEs, however we created a reference pME catalog from the high-confidence structural variants sets reported by the genome in a bottle consortium (GIAB), which offer an exhaustive representation of the SVs > 50bp found in the genome of HG002/NA24385 compared to the reference hs37h5 (GRCh37/hg19, accessible at: https://ftp-trace.ncbi.nlm.nih.gov/ReferenceSamples/giab/release/references/GRCh37/). The GIAB/HG002 variant calls rely on the combination of 4 sequencing technologies (Illumina 250bp paired-end, Illumina mate-pair, 10X genomics linked-reads, and PacBio HiFi) and 19 SV callers (accessible at: https://ftp-trace.ncbi.nlm.nih.gov/ReferenceSamples/giab/data/AshkenazimTrio/analysis/NIST_S <u>Vs_Integration_v0.6/</u>). To retain SVs representing pMEs, we applied the GraffiTE annotation step on the HG002_SVs_Tier1_v0.6.vcf.gz VCF file using the --vcf input option. We collected the complete list of human TE consensus sequences from DFAM v.3.6^91^ in fasta format using the FamDB toolkit (available online at: https://github.com/Dfam-consortium/FamDB). We then retained pMEs with a single RepeatMasker hit along more then 80% of the sequence and annotated as Alu, LINE-1 or SVA, with a minimum size of 250 bp. This file is accessible online (GIAB.Tier1.Alu.L1.SVA.250bp.vcf) at: https://github.com/cgroza/GraffiTE/tree/main/paper#giabhg002-benchmark). To test GraffiTE and compare its outputs with other existing tools (see below), The diploid assembly used in this benchmark was produced by the Telomere-to-Telomere consortium^92^ and available online at: https://github.com/marbl/HG002. We also gathered ∼50X of Illumina 2x250bp short-reads available for HG002 (140528_D00360_0018_AH8VC6ADXX and 140528_D00360_0019_BH8VDAADXX/ accessible at https://ftp-trace.ncbi.nlm.nih.gov/giab/ftp/data/AshkenazimTrio/HG002_NA24385_son/NIST_HiS <u>eq_HG002_Homogeneity-10953946/HG002_HiSeq300x_fastq/</u>) and ∼32X of PacBio high-fidelity (HiFi) long-reads (accessible at: https://ftp-trace.ncbi.nlm.nih.gov/ReferenceSamples/giab/data/AshkenazimTrio/HG002_NA2438 <u>5_son/PacBio_SequelII_CCS_11kb/reads/</u>). The read sets (long and short) were subsequently re-sampled to 5, 10, 20 and 30X coverage using Rasusa^93^.

### pME detection evaluation

In order to assess the performance of GraffiTE on pME detection, we ran the pipeline with different combinations of tools and data (Table S1). First, we ran GraffiTE using only the maternal and paternal assemblies as input; in this condition (further labeled GT-sv), SVIM-asm is the only software used to detect variants. Alternatively, we ran GraffiTE on long-read data only, using either 5, 10, 20 or 30X coverage. This analysis implied the use of Sniffles 2^52^ and is further referred to as GT-sn. Finally, we also ran GraffiTE with a combination of alternative assemblies and increasing coverage of long-reads (GT-sv-sn). GraffiTE was run with the flag --mammal, which identifies L1 with 5’ inversion as a single pME and annotates SVA VNTR polymorphism (SVA-VNTR polymorphisms were further discarded for the benchmark). We filtered the GraffiTE output VCFs using the same criteria that for the GIAB reference call: we retained only variants with a single RepeatMasker hit to an Alu, L1 or SVA element and a variant length >= 250bp. According to their input specification, we also ran a collection of representative methods either using long-reads (TLDR^28^) or short-reads (MELT2^6^, MEGAnE^68^). Details about the parameters used and output conversion in the VCF format can be found in Supplementary Methods 1b. To asses pME search performances, comparison between the different methods and the constructed GIAB pME reference set were carried out using the R package sveval^69^ with defaults parameters and provided the file HG002_SVs_Tier1_v0.6.bed (https://ftp-trace.ncbi.nlm.nih.gov/ReferenceSamples/giab/data/AshkenazimTrio/analysis/NIST_ <u>SVs_Integration_v0.6/HG002_SVs_Tier1_v0.6.bed</u>) which “*should contain close to 100 % of true insertions and deletions >=50bp*’’ according to the GIAB documentation.

### pME genotyping evaluation

We further sought to evaluate the performance of the graph-genotyping tools implemented in GraffiTE, specifically Pangenie (for short-read data) and GraphAligner/vg call (for long-read data). We selected the pME annotated VCFs from GraffiTE for the runs GT-sv, GT-sn and GT-sv-sn and performed graph-genotyping with short- or long-reads at 30X coverage (Table S1). We also used the results from MELT2 and MEGAnE (at 30X coverage) for evaluation, as these tools provide bi-allelic genotypes for the detected pMEs. Genotyping performance was evaluated using sveval with the additional parameters: geno.eval=TRUE, method="bipartite", stitch.hets=TRUE, merge.hets=FALSE.

### Application Examples

#### Human pangenome

We ran GraffiTE on the 47 HPRC diploid assemblies^34^ (94 inputs as haploid genomes) using the human transposable element models from DFAM 3.6^91^ and the hg38 reference genome^94^. The raw output of GraffiTE (VCF file) was then filtered to retain Alu, L1 and SVA elements using the same criterias as for the variants of the GIAB benchmark. We then calculated discovery curves, separating polymorphisms into rare (<=5% frequency), common (between 5% and 95%) and “fixed” (>95%) categories, permuting the order of genomes 100 times. We recapitulated the expected population structure by running principal coordinate analysis on pME presence/absence calls present in the GraffiTE pangenome.vcf output (no graph-genotyping was performed as only assemblies were used) with the ade4 R package^95^ (distance = coefficient S5 of ^96^) using pMEs with a pangenome frequency between 5% and 95%. Samples’ metadata were obtained from^34^. We further performed a comparison of the total number of pMEs discovered between the results of GraffiTE using minimap2/svim-asm as SV search tool and the reference-free methods used in^71^ based on the Minigraph/Cactus software. We converted the genome graph produced by the authors (GFA) to VCF using vg deconstruct. We then used the VCF file as input for GraffiTE (--vcf argument). Upon analysis, we filtered the VCF as described above to retain Alu, L1 and SVA. Before comparing the with the earlier VCF (obtained with GraffiTE using minimap2/svim-asm on the assemblies), we removed duplicated SVs in the Minigraph/Cactus VCF (see Supplementary Methods).

#### Drosophila melanogaster pangenome

We obtained genome assemblies and raw long read sets for 30 genomes resequenced by Rech et al.^8^ and using ONT long-reads technology (details can be found on table S2 of Rech et al.^8^, for genome with “technology: ONT”). We then applied GraffiTE, using for each sample both the assembly and long-reads (ONT) set for SV search, and GraphAligner/vg call to genotype pMEs in the TE pangenome with long-reads (GT-sv-sn-GA mode, see table S1-B). To annotate pMEs, we used the new “MCTE” library produced by Rech et al.^8^. The final GraffiTE VCF was filtered to retain variants with a single TE hit (n_hits=1) and fixed variants relative to dm6 after genotyping were removed, as well as variants with missing genotypes. We calculated discovery curves for pMEs detected in the *D. melanogaster* pangenome for low frequency (N = 1/30 to 2/30 genomes), common (3/30 ≤ N ≤ 28/30 genomes) and fixed (N = 29/30 to 30/30 genomes), permuting and sampling genomes 100 times for each frequency bin.

#### Zea mays bz *locus*

We applied GraffiTE to 23 long-read sets from 23 cultivars published by Hufford et al. (2021; doi:10.1126/science.abg5289) using the telomere-to-telomere (Zm-Mo17-T2T ) reference of Mo17^75^ and the TE library NAM.EDTA2.0.0.MTEC02052020.TElib.fa used in^66^ and available at https://github.com/oushujun/PopTEvo. We chose Zm-Mo17-T2T as reference as it is expected to be the most complete assembly available for a maize strain. We used the VCF file produced by GraffiTE to generate a graph genome (in GFA format) at a resolution of 32 bp (node length), and drew its layout with Bandage^76^. Next, we mapped the TE annotations from GraffiTE to the graph genome (Supplementary Methods 2c) and further annotated the shared paths of the graph using RepeatMasker and the NAM.EDTA2.0.0.MTEC02052020.TElib.fa library to produce an annotation file (CSV) to load with Bandage. Additionally, we obtained 8 haplotypes analyzed by^77^ and mapped these haplotypes to the graph in order to highlight the path of these particular varieties relative to the different polymorphisms detected by GraffiTE. We first merged all possible nodes using the dedicated Bandage functionality and mapped the haplotype using blastn (-word_size 22) using the onboard blastn tool.

#### Cannabis sativa *pangenome*

We used the cs10 reference genome (GenBank: GCA_900626175.2) and 9 publicly available alternative assemblies (Table S3) to build a TE-graph genome with GraffiTE. First, we created a *de-novo* TE library using RepeatModeler2^84^ and clustered 1737 models automatically generated with 128 models present in Repbase v.27.06^97^ to reduce redundancy and remove the models already present in Repbase from the automatic library (Supplementary Methods 2.b). We then applied GraffiTE to the 9 alternative assemblies, using cs10 as reference and the generated TE library. Conversely to *D. melanogaster* and human samples, pMEs for *C. sativa* were previously undocumented. After running GraffiTE, we noticed that several reported pMEs had annotations composed of a patchwork of hits against different consensus sequences of the library (which is common with non-curated libraries). To circumvent this, and demonstrate the use of GraffiTE with non-model organisms, we further clustered the sequences of all candidate pMEs (variants reported in the pangenome.vcf output) among the 9 genomes using MMseqs2^98^ (parameters easy-cluster --cov-mode 0 -c 0.8 --min-seq-id 0.8 -s 100 --threads 8 --exact-kmer-matching 1, see also Figure 6A). This clustering allowed to group together pME loci which share at least 80% of their sequence with a minimum identity of 80%. We then retained clusters with a minimum of 3 loci as putative TE families and with a size of 200 to 40,000 bp. The top 100 (in number of pME loci per cluster/family across the *C. sativa* pangenome) were analyzed and representative sequences of 4 families were manually annotated to illustrate the diversity of pME families in *C. sativa*. The analysis was replicated adding as input 5 publicly available long read sets supporting 5 out of the 9 assemblies used (see Supplementary Methods 2d).

## Acknowledgments

We are grateful to Casey Bergman who provided insightful comments on the manuscript. We also thank Clémentine Vitte for sharing her expert knowledge about TE variation in maize.

## Author Contributions

**CR.G.** Developed the software, ran experiments, performed analyses and wrote the manuscript. **CL.G.** Designed and developed the software, designed and ran experiments, performed analyses and wrote the manuscript. **X.C.** performed software testing and ran the short-reads experiments. **T.J.W.** and **G.B.** supervised the work. All authors contributed to the manuscript.

## Data availability

Source data are provided with this paper (pMEs discovery and genotyping VCFs and formatted table for simulations, GIAB benchmark results, application examples), as well as the scripts used for analysis are available in the Zenodo repository https://zenodo.org/doi/10.5281/zenodo.11391566 and our Github page: https://github.com/cgroza/GraffiTE/tree/main/paper

## Code availability

GraffiTE is open source (MIT License). The source code and complete manual for GraffiTE are freely available on Github: https://github.com/cgroza/GraffiTE. A frozen version of the Github repository, generated on June 26th of 2024 is available at: https://doi.org/10.5281/zenodo.12538787.

