## Supplementary Methods for "A Unified Framework to Analyze Transposable Element Insertion Polymorphisms using Graph Genomes"

#### SUPPLEMENTARY INFORMATION

**Authors**: Cristian Groza, Xun Chen, Travis J. Wheeler, Guillaume Bourque and Clément Goubert

Supplementary Methods
Supplementary Figures
Supplementary Tables

### Supplementary Methods

#### 1. Evaluation of GraffiTE performance

##### a. Simulations

i. General model

In order to evaluate the performance of GraffiTE, we performed pMEs simulations based on two distinct model species: one in human and one in maize. The details for each model are reported below. Briefly, the simulation process consists of generating pairs of genomes: one used as reference, and one used as an alternative sample. For each simulation, the alternative sample/genome contains pME insertions or deletions relative to the reference one. Each pME is simulated with target site duplications (TSD) whose sizes are variable depending on the TE family considered (see below). The pMEs simulated can also be either a full-length copy or a fragment; the inserted or deleted sequences are taken from the set of reference TE annotation provided for each species (see below). Our simulator generates a VCF file with 20 pMEs (true positives) and 20 background SVs (false positives) over random stretches of 10Mbp using the human chromosome 22 or the maize chromosome 10 as starting point. Next, the VCF and corresponding 10Mb genomic region are given to SimuG[1] to produce a new genome based on the simulated variants. The simulation software includes customisable parameters such as the number of variants to simulate and the source/target chromosome and is available at https://github.com/clemgoub/pMEsim. In addition, putative false positives are generated by modeling background insertion or deletions. The sequences and breakpoints of these background SVs are randomly chosen, avoiding “N” insertion sites, intervals with more than 10% “N” residues as well as intervals overlapping with the 20 previously simulated pMEs. Background SV sizes are randomly picked from the catalog of structural variants (insertion or deletions) available for each model (see below). 100 Simulations were performed per species (100 pairs of reference and alternate genomes). Since our goal was to test the ability of the tools to call presence/absence, we did not simulate heterozygosity in the reads, thus simulated genomes were haploid. Next, we generate two read-sets per simulated genome: one using the ONT error model (Oxford Nanopore) and one using the Sequel II error model (PacBio CLR reads). Sequel II / CLR reads are simulated with 10 passes and  a target  coverage of 15.23 X with human data and 11.04 X with maize data (instead of 10X), in order to yield 10X after converting them into hi-fidelity using ccs (Pacific Bioscience, v.6.4.0). For each simulation, we ran GraffiTE with either of the five following modes: assemblies only (GT-sv: svim-asm), long-read only (GT-sn: Sniffles2, ONT), long-reads only (GT-sn: Sniffles2, hifi) and assembly+long-reads (GT-sv-sn: svim-asm + Sniffles2 ONT), and assembly+long-reads (GT-sv-sn: svim-asm + Sniffles2 hifi).

ii. Human model:

In order to simulate realistic pMEs in the human genome, AluY copies are required to be at least 250 bp long and L1/SVA copies >= 900 bp. The pool of pME to insert or remove from the chromosome 22 was obtained from the RepeatMasker track provided by UCSC for hg38 (https://hgdownload.soe.ucsc.edu/goldenPath/hg38/bigZips/hg38.fa.out.gz), and curated to retain AluY, L1 (L1HS, L1PA2) and SVA (SVA_E-F). For each simulation, 14 insertions and 6 deletions are performed. Furthermore, Alu, L1 and SVA were picked with the ratio 0.7/0.2/0.1 to mimic known pME proportions[2,3]. The ratio used slightly over-represents deletions and rarer pMEs such as L1 and SVA in order to obtain enough replicates in the statistical analyses. TSD length for Alu, L1 and SVA were taken within a range of 4 (min TSD length distinguishable by GraffiTE) and 22 as described in[4,5]. Background SV sizes are simulated using the length distribution reported by Zook et al.[6] for the individual HG002.

iii. Maize model:

Maize simulations were done using the same framework as the human model. We selected TE families from two major clades active in the maize genome: Class II DNA/TIR (DTA) of the *hAT* superfamily and Class I LTR retrotransposons of the superfamilies *Copia* and *Ty3-Metaviridae* (formerly known as *Gypsy*). The selected families and corresponding classification are given below.

ansuya_AC207803_9728 Gypsy
chr5_P_133655019 Copia
chr6_P_87978263 Copia
cinful_zeon_AC177930_407 Gypsy
cinful_zeon_AC208228_9805 Gypsy

| | |
|---|---|
| cinful_zeon_AC209373_10201 | Gypsy |
| DTA_ZM00106_consensus | DTA |
| DTA_ZM00135_consensus | DTA |
| DTA_ZM00147_consensus | DTA |
| DTA_ZM00170_consensus | DTA |
| DTA_ZM00172_consensus | DTA |
| DTA_ZM00244_consensus | DTA |
| DTA_ZM00254_consensus | DTA |
| DTA_ZM00275_consensus | DTA |
| DTA_ZM00342_consensus | DTA |
| flip_AC194904_4319 | Gypsy |
| gunu_AC209758_10359 | Copia |
| huck_AC186577_1525 | Gypsy |
| huck_AC186603_1556 | Gypsy |
| huck_AC190900_2713 | Gypsy |
| huck_AC191259_3001 | Gypsy |
| huck_AC193313_3542 | Gypsy |
| huck_AC194973_4393 | Gypsy |
| huck_AC199418_6452 | Gypsy |
| huck_AC199444_6460 | Gypsy |
| huck_AC203007_7610 | Gypsy |
| huck_AC210079_10574 | Gypsy |
| huck_AC212331_11708 | Gypsy |
| huck_AC213042_12069 | Gypsy |
| huck_AC216048_13250 | Gypsy |
| ji_AC193479_3665 | Copia |
| ji_AC200613_6936 | Copia |
| ji_AC202456_7443 | Copia |
| nida_AC206942_9401 | Copia |
| opie_AC187149_1780 | Copia |
| opie_AC188002_2029 | Copia |
| opie_AC196469_5133 | Copia |
| opie_AC197084_5382 | Copia |
| opie_AC197201_5474 | Copia |

| | |
|---|---|
| opie_AC197691_5727 | Copia |
| opie_AC198924_6206 | Copia |
| opie_AC202020_7258 | Copia |
| opie_AC202033_7274 | Copia |
| opie_AC214122_12495 | Copia |
| opie_AC217577_13524 | Copia |
| TE_00000045 | DTA |
| TE_00000312 | DTA |
| TE_00000564 | DTA |
| TE_00000685 | DTA |
| TE_00001225 | DTA |
| TE_00001660 | DTA |
| TE_00001940 | DTA |
| TE_00002079 | DTA |
| TE_00003428_INT | Gypsy |
| TE_00003751_INT | Gypsy |
| TE_00006785 | DTA |
| TE_00007667 | DTA |
| TE_00007995 | DTA |
| TE_00013629_INT | Gypsy |
| TE_00013713_INT | Gypsy |

These families were selected after analyzing the TE annotations for the assembly Zm-Mo17-REFERENCE-CAU-2.0.fa[7] available at: https://data.cyverse.org/dav-anon/iplant/home/laijs/Zm-Mo17-REFERENCE-CAU-2.0/. We request that the selected families have at least 10 copies >= 99% of the length of the consensus sequence (full-length TE) and less than 10% divergence to the consensus on average in order to favor likely active TEs. Chromosome 10 present in Zm-Mo17-REFERENCE-CAU-2.0.fa was used as the template for the simulation. For DTA elements, TSDs of 8bp were simulated while LTR/Copia and LTR/Ty3 elements were simulated with TSDs ranging between 4 and 6bp based on[8]. Background SV length (false positives) were simulated using the SV length distributions reported by[9].

##### b. Comparison to existing methods

*TLDR (long-reads)*

We applied TLDR[10] (accessed online at https://github.com/adamewing/tldr, April 2023) on the long-read alignments produced by GraffiTE with minimap2. TLDR was run with default parameters and the DFAM 3.6 human TE library. Outputs with a "PASS" flag were converted into VCF using the rounded mean of column 3 and 4 (start end for the breakpoint) of the native output table for the POS field in the VCF (this slightly improved TLDR F1 scores compared to using column 3 "start" as POS field). TLDR only reports non-reference pMEs (INS).

*MELT2, MEGAnE (short-reads only)*

We used the short-read sets generated from HG002 at 5, 10, 20 and 30X coverage to detect reference (DEL) and non-reference (INS) pMEs. Both MELT2 (version: v2.2.2) and MEGAnE (version: v1.0.0) were applied with default parameters. Identified polymorphic Alu, LINE1 and SVA elements were used for the comparative analysis. We found that filtering MELT2 and MEGAnE for the "PASS" flag did increase significantly their false negative rate, and thus did include all the variants reported in order to maximize their F1 score in the comparison

In order to match GraffiTE filters, we retained Alu, L1 and SVA calls >= 250bp in each output dataset.

#### 2. Application Examples

##### a. Human HPRC pangenome

Besides the analysis presented in Figure 3, we also compared the GraffiTE results using minimap2/svim-asm to the data present in the genome graph (GFA) produced by[11]. After conversion to VCF, the data were filtered to remove duplicated SVs. These were caused by SNP or indels observed between assemblies however for variants sharing the same length and VCF coordinates. We observed that these are likely due to sequencing errors, in particular those in homopolymers which are present as poly-A tails for human TEs. In addition, these apparent duplications share the same TSD sequence; making it unlikely, in humans, that two TEs from the same family will insert at the same locus.

##### b. *Drosophila* pangenome

In their original publication, Rech et al.[12] reported pMEs by comparing TE present in each assembly, as annotated by the pipeline REPET, with those given for the reference genome ISO-1/dm6. Putative pMEs are then reported in a bed file whose coordinates are relative to each genome; using an ad-hoc script, the authors further converted these coordinates to the reference genome. This approach (annotate first, detect variant next) is orthogonal to GraffiTE (detect variants first, annotate them next) but proved to be challenging to convert back into the standard VCF format with confidence. pME calls from Rech et al. (Table S10) were initially converted into a VCF file to compare results with GraffiTE. The conversion considered an insertion (INS) if "ISO-1" (reference genome) was absent from the "genome" column (meaning the TE is absent in the reference), and labeled deletion (DEL) loci for which ISO-1/dm6 is reported in the "genome" column. For INS, the variant length could be variable between members of the pangenome, and was thus reported in VCF format by calculating the median length from the individual values in table S10. For DEL, the length was taken as reported for the reference genome. For some DEL loci, the TE length was either longer or shorter than reported for the reference, making them inconsistent with the given DEL classification; thus we systematically discarded loci with +/- 10% length difference between individual genome and the reference. We recalculated the pME frequency among genomes in this subset, removing loci fixed between the 30 genomes and ISO-1/dm6, since GraffiTE would not be able to detect them.

Upon comparison of the generated VCF for Rech et al.[12], we noticed a seemingly poor overlap with our data: sveval (see Methods) showed that 10,252 pME loci reported by GraffiTE could be matched to 10,253 loci of Rech et al., representing half and ~2/3 of each dataset (restricted to euchromatic regions), respectively. In order to understand the reasons behind the observed discrepancies, we randomly pulled 96 pME insertions predicted by Rech et al. (2022), and for which GraffiTE seemed to fail to produce a matching locus according to sveval. For each pME, we manually searched for the closest two variants reported in the GraffiTE output. This search revealed that 47.9% of the Rech loci (45 pME) could actually be matched unambiguously to a GraffiTE locus ("MATCH" category). The reasons for not being reported by sveval are multiple, and include for this category a single of either (i) a distance > 30bp between the variant breakpoints, (ii) significant difference in length between the variants reported, or (iii) multiple TEs hits reported within a GraffiTE variant (such variants were filtered out from the final GraffiTE

VCF). Furthermore, 8 pMEs were labeled as “possible MATCH”, when two or more of these factors contributed to the discrepancies. Another 4 loci (4.3%) were initially discovered by the SV callers implemented in GraffiTE; however, as their sequences did not contain a RepeatMasker hit against a known TE for at least 80% of their sequence, these variants were rejected at the annotation/filtering step of the pipeline. Finally, 37 pMEs could not be matched at all and were further examined visually in order to assess whether or not the variant predicted by Rech et al.[12] was likely a true or a false positive. For each of them, we extracted the genome region containing the pME +/- 2kb of flanking sequences from the assembly from which the pME was predicted (if the pME was reported in multiple assemblies, only the first genome reported was used to extract the locus in question). The 5’ and 3’ flanks were also extracted alone. Then, for each insertion, 3 sequences (“full” including the TE +/- 2kb of flanking, the 5’ and 3’ flanks) were mapped to the reference genome dm6 with the online implementation of blat, using the ucsc web server (https://genome.ucsc.edu/cgi-bin/hgGateway?hgsid=1667170868_JpFqs9rTDHR1OnaApFiU2BUHvbuR). A true positive was expected to show the 5’ and 3’ flanks mapping next to each other on either side of the predicted breakpoint. In some instances, blat can also map the “full” sequence displaying the TE as insertion (see Supplementary Figures 4-10). We considered false positives when the mapping did show that the predicted pME was in fact already in the dm6 reference, or if proper mapping of the flanking regions at the expected location could not be confirmed. Accordingly, 22 pME (23.4%) were classified as false positives, and 15 (16%) were found to be true positives.
Based on these exploratory results, we concluded that an automatic conversion of the Rech et al.[12] data to the VCF format introduced too many ambiguities to perform a systematic comparison. Furthermore, the absence of an independent truth set (e.g., genotyping from PCR, or alternative predictions from a benchmarked third-party software) prevents us from objectively evaluating the respective performances of each method (we do not consider the manual observation described above objective enough to contribute to the main analysis).

##### c. Maize pangenome

We ran GraffiTE on the Zea mays genome using the Zm-Mo17-T2T as a reference genome[7]. We discovered TEs with GraffiTE from the following assemblies: Zm-Oh43, Zm-Ms71, Zm-Il14H, Zm-Ki3, Zm-CML228, Zm-Tx303, Zm-Mo18W, Zm-NC358, Zm-Tzi8, Zm-CML103, Zm-Oh7B, Zm-CML247, Zm-CML322, Zm-NC350, Zm-CML277, Zm-M162W, Zm-HP301,

Zm-B97, Zm-CML69, Zm-Ki11, Zm-B73, Zm-CML333, Zm-P39, Zm-M37W, Zm-Ky21, Zm-CML52, Zm-A188[13]. To these assemblies, we also added contigs describing the *bz* locus from[14]**.** In following experiment, we also discovered TEs from the following PacBio long-read SRA datasets: SAMEA5313298, SAMEA5313299, SAMEA5313300, SAMEA5313301, SAMEA5313302, SAMEA5313303, SAMEA5313304, SAMEA5313305, SAMEA5313306, SAMEA5313307, SAMEA5313308, SAMEA5313309, SAMEA5313310, SAMEA5313311, SAMEA5313312, SAMEA5313313, SAMEA5313314, SAMEA5313315, SAMEA5313316, SAMEA5313318, SAMEA5313319, SAMEA5313320, SAMEA5313321, SAMEA5313322. In all cases, we used the TE library available at: https://download.maizegdb.org/Transposable_elements/NAM.EDTA2.0.0.MTEC02052020.TElib.fa.gz to identify transposable elements in maize variants.

We used the following command to run the *Zea mays* assemblies:

```
nextflow run GraffiTE/main.nf --assemblies assemblies.csv
--TE_library NAM.EDTA2.0.0.MTEC02052020.TElib.fa --reference
GCA_022117705.1_Zm-Mo17-REFERENCE-CAU-T2T-assembly_genomic.fna
--genotype false -with-singularity graffite_latest.sif
```

For running the *Zea mays* long-reads, we used:

```
nextflow run GraffiTE/main.nf --longreads longreads.csv --TE_library
NAM.EDTA2.0.0.MTEC02052020.TElib.fa --reference
GCA_022117705.1_Zm-Mo17-REFERENCE-CAU-T2T-assembly_genomic.fna
--graph_method graphaligner -with-singularity graffite_latest.sif
```

Code to repeat mask and annotate the *bz* locus subgraph is available in the github repository under `paper/Applications_Examples/Zea_mays`. In summary, we extract the *bz* locus subgraph, chunking the nodes in 32 bp pieces using `vg chunk`. Then, we extract the sequences and alignments of all the paths that run through this graph using `vg paths`, obtaining a FASTA and a GAF file of all reference and SV sequences and alignments. We run `RepeatMasker -s -nolow -lib NAM.EDTA2.0.0.MTEC02052020.TElib.fa` on these sequences to obtain the repeat annotation, then join this information to the nodes in the graph using `join_annotation.R`. The outputs may then be loaded and visualized with Bandage.

**d. *Cannabis sativa* Pan-genome**

Automatically generated TE models (consensus sequences) from RepeatModeler 2 were clustered with *C. sativa* sequences gathered from Repbase using MMseqs2 [15] with the parameters "easy-cluster --cov-mode 1 -c 0.8 --min-seq-id 0.8 -s 100 --exact-kmer-matching 1" and prioritized manually curated models from Repbase over automatic RepeatMasker2 consensus as representative of each cluster for a final number of models of 1680. The final library, minus the Repbase models, was submitted to the open-source repository DFAM [16] as a non-curated library. We performed an additional analysis, replicating the experiment described in Methods, with the addition of 5 long read sets (GT-svsn mode). Each additional read (SRR10189115 through SRR10189120, SRR12825097, SRR21901303, SRR7274799 and SRR7274947) set correspond to the raw sequencing data supporting the assemblies JL (GCA_013030365.1), Cannbio-2 (GCA_016165845.1), Pink Pepper (GCA_029168945.1), Finola (GCA_003417725.2) and Purple Kush (GCA_000230575.5)

### Supplementary Figures

- **Supplementary Figure 1:** Pipeline workflow
- **Supplementary Figure 2:** TSD module
- **Supplementary Figure 3:** Details of the RepeatMasker annotations characteristics of canonical and twin-primed LINE1
- **Supplementary Figure 4:** Comparison of pMEs counts according to the SV discovery method for the HPRC pangenome.
- **Supplementary Figures 5-8:** Examples of false positives in Rech et al., 2022
- **Supplementary Figures 9-11:** Examples of true positives in Rech et al., 2022
- **Supplementary Figure 12:** Comparison between GT-svsn-GA (long reads genotyping) and GT-svsn-PA (short reads genotyping) for *D. melanogaster* application example.
- **Supplementary Figure 13:** Comparison of the total number of candidate pME in the *C. sativa* pangenome whether assemblies (GT-sv) or long-reads (GT-sn) are used for pME search.

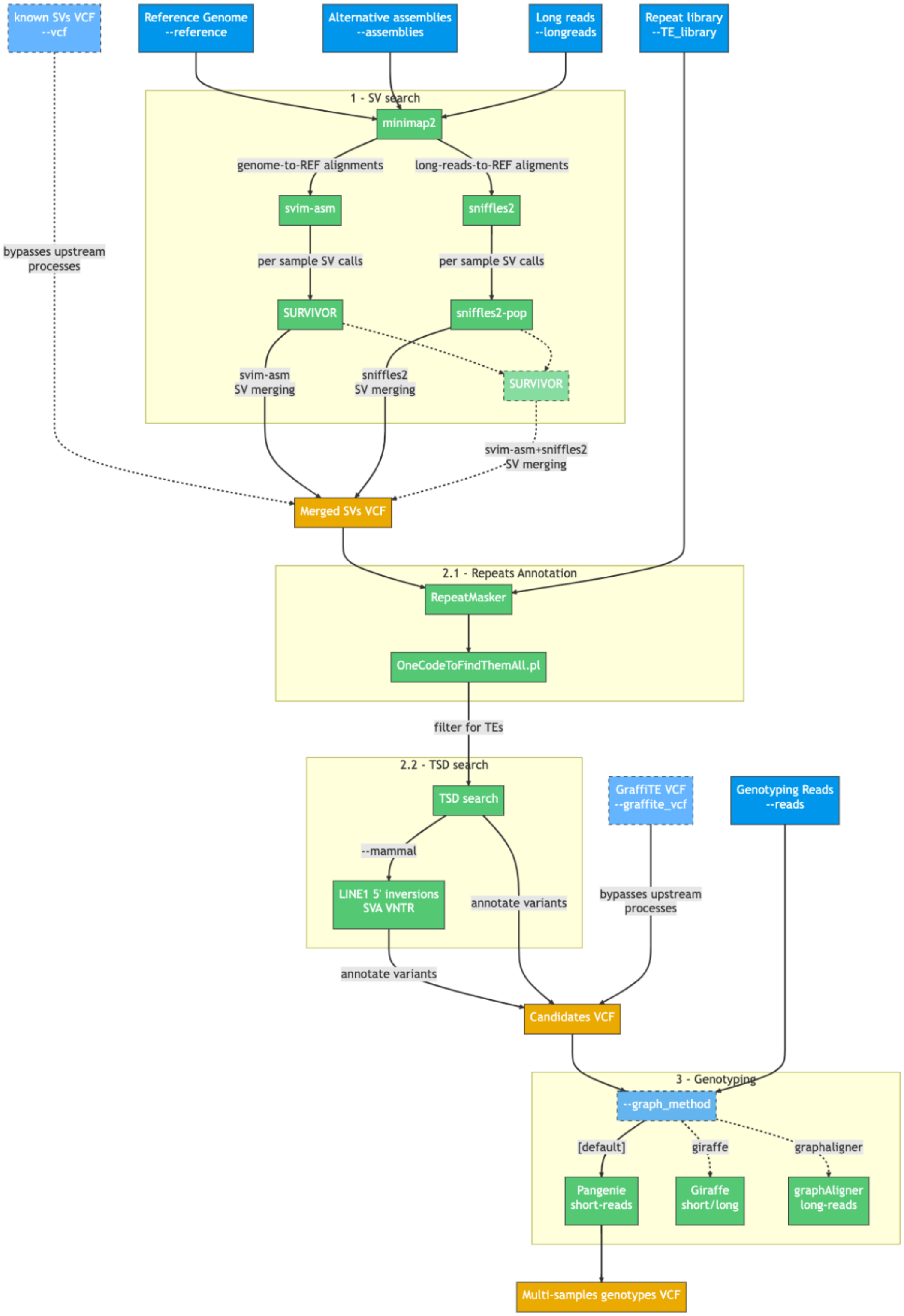


**Supplementary Figure 1: Detailed GraffiTE workflow**. Blue boxes indicate user inputs with the associated GraffiTE parameters. Faded blue boxes depict shortcuts or options in the pipeline. Green boxes represent the main software and scripts implemented. Orange boxes highlight the main VCF outputs of GraffiTE. Dotted lines represent alternative/optional paths in the workflow.

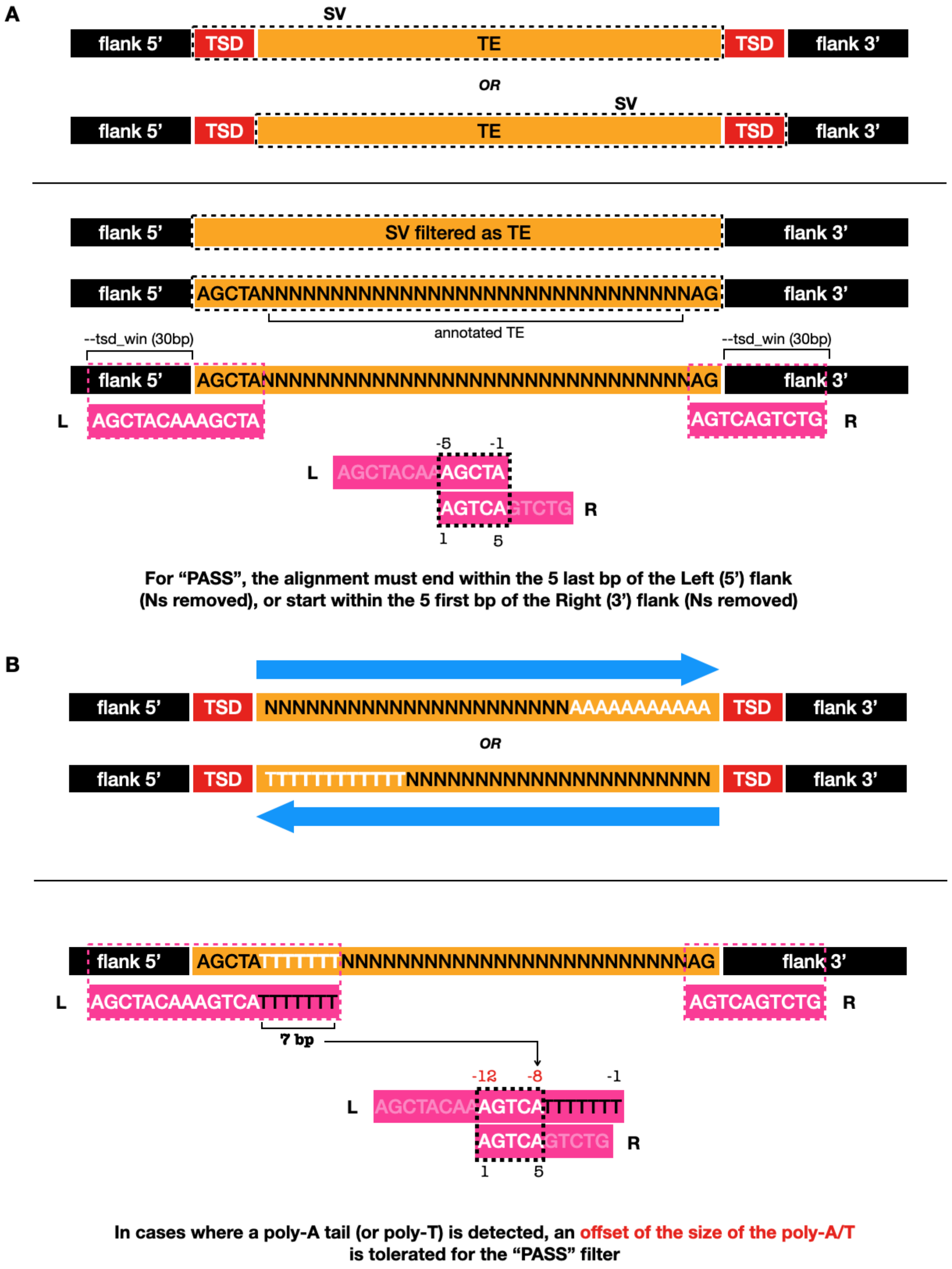
A
SV
flank 5'
TSD
TE
TSD
flank 3'
OR
SV
flank 5'
TSD
TE
TSD
flank 3'
flank 5'
SV filtered as TE
flank 3'
flank 5'
AGCTANNNNNNNNNNNNNNNNNNNNNNNNNNNNNNNNNNNNAG
flank 3'
annotated TE
--tsd_win (30bp)
--tsd_win (30bp)
flank 5'
AGCTANNNNNNNNNNNNNNNNNNNNNNNNNNNNNNNNNNNNAG
flank 3'
L
AGCTACAAAGCTA
AGTCAGTCTG
R
-5
-1
L
AGCTACAAAGCTA
AGTCAGTCTG
R
1
5
For "PASS", the alignment must end within the 5 last bp of the Left (5') flank (Ns removed), or start within the 5 first bp of the Right (3') flank (Ns removed)
B
flank 5'
TSD
NNNNNNNNNNNNNNNNNNNNNNNNAAAAAAAAAAA
TSD
flank 3'
OR
flank 5'
TSD
TTTTTTTTTTTTNNNNNNNNNNNNNNNNNNNNNNNN
TSD
flank 3'
flank 5'
AGCTATTTTTTTNNNNNNNNNNNNNNNNNNNNNNNNNNNNNAG
flank 3'
L
AGCTACAAAGTCATTTTTTT
AGTCAGTCTG
R
7 bp
-12
-8
-1
L
AGCTACAAAGTCATTTTTTT
AGTCAGTCTG
R
1
5
In cases where a poly-A tail (or poly-T) is detected, an offset of the size of the poly-A/T is tolerated for the "PASS" filter

**Supplementary Figure 2: TSD search module. A**: TSDs are searched by comparing >=30bp of the flanking regions of an annotated TE within each reported SV (N's stretches). An empirical filter gives a "PASS" flag (and reports the TSD in the output VCF) if the TSD alignments starts within the 5 bp flanking the TE annotation (other matches are reported in the dedicated TSD output tables). **B**: In case a poly-A (or T) tail is detected, an offset of the poly-A(T) size is tolerated to receive a "PASS" flag.

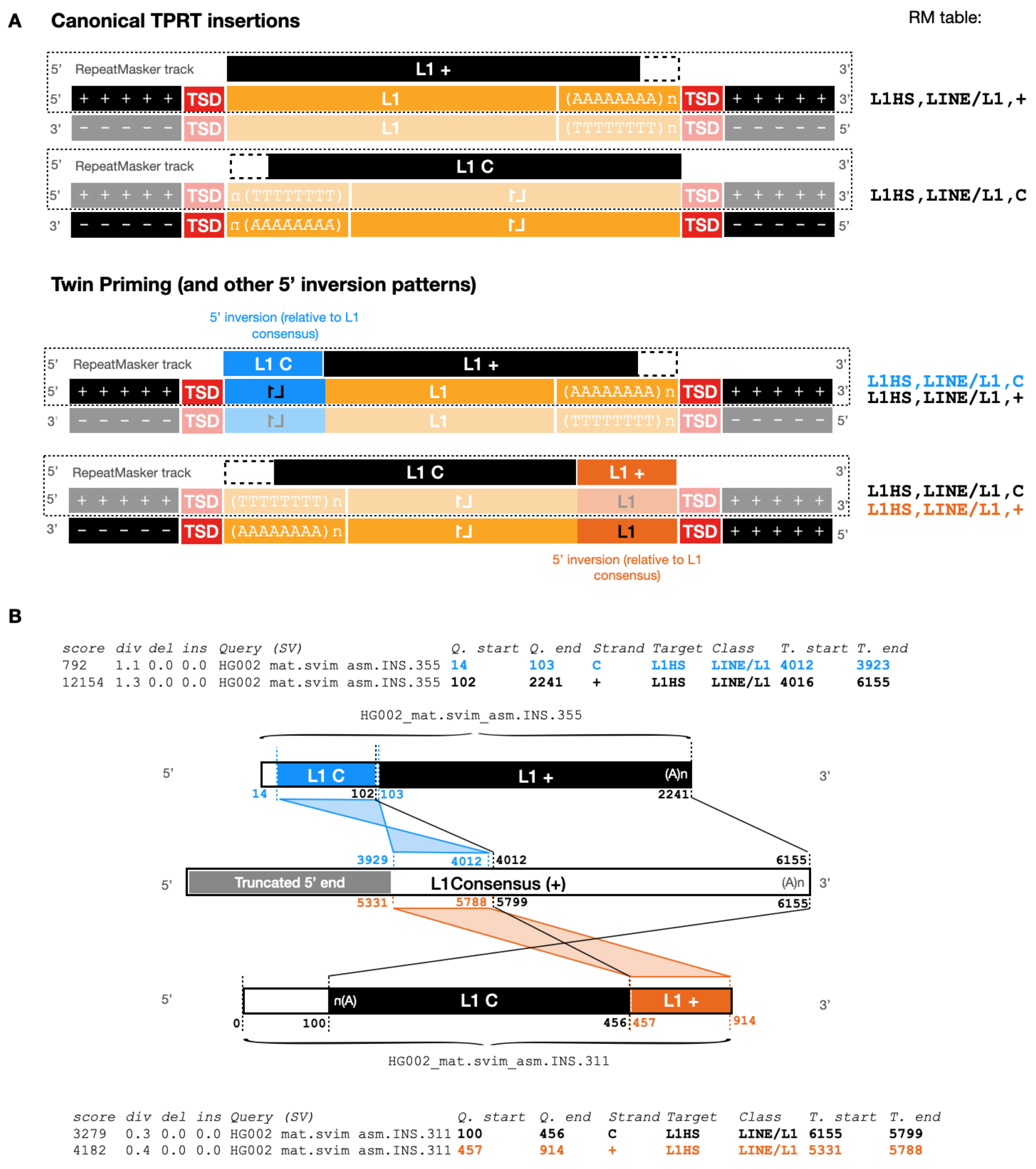


**Supplementary Figure 3: Details of the RepeatMasker annotations characteristics of canonical and twin-primed LINE1** (as well as other mechanisms leading to 5' inversion of a LINE1 copy). **A**: comparison of canonical and 5' inverted LINE1 insertions, with corresponding RepeatMasker annotation (right). **B**: Details of the clues used to determine the orientation (+ or -) of a LINE1 copy with 5' inversion based on the RepeatMasker outputs.

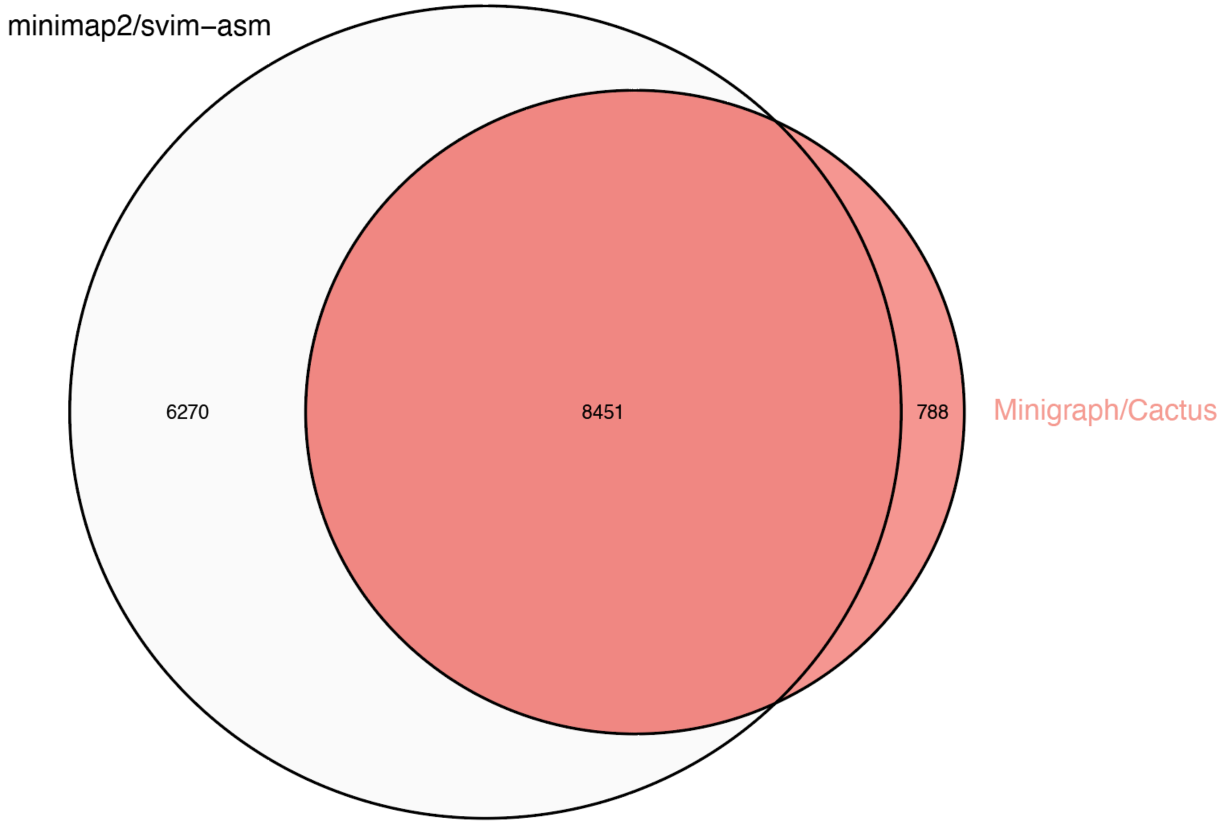


**Supplementary Figure 4. Comparison of pMEs counts according to the SV discovery method for the HPRC pangenome.** pMEs comparison is made between the default input for GraffiTE (minimap2/svim-asm, as presented in the main text) and the VCF conversion of the GFA produced by (Minigraph/Cactus). The Minigraph/Cactus set was also filtered to keep 1 SV entry per unique locus (SNPs and indels detected between samples at the same SV resulted in independent entries in the Minigraph/Cactus dataset). For each dataset (VCF file) only variants with a single hit to a Alu, L1 and SVA element are kept (see main text).

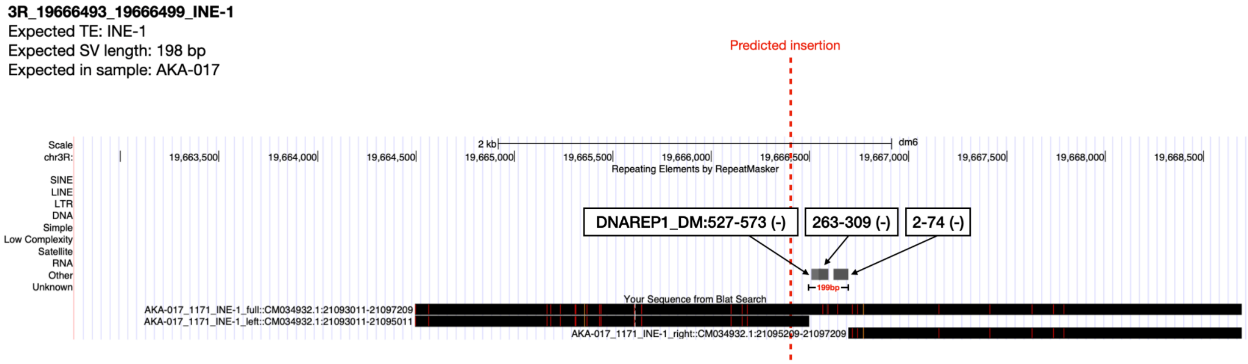


**Supplementary Figure 5. False positive insertion in the dm6 assembly reported by Rech et al., 2022.** Screenshot from the UCSC genome browser showing Blat mapping of the predicted insertion sequence +/- 2kb of flanking. Three sequences are mapped: full insertion with flanking ("full"), left and right flanking only ("left" and "right"). The predicted insertion sequence from the genome AKA-017 fully maps onto the reference dm6. The predicted INE-1 TE is annotated as DNAREP1_DM in multiple fragments in the UCSC representation of the dm6 assembly.

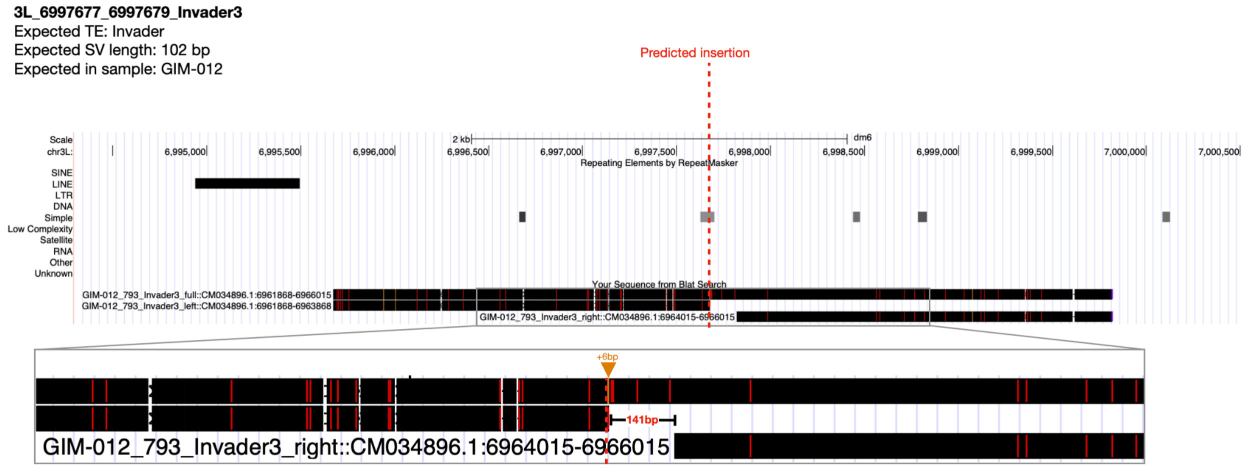


**Supplementary Figure 6. False positive insertion in the dm6 assembly reported by Rech et al., 2022.** Screenshot from the UCSC genome browser showing Blat mapping of the predicted insertion sequence +/- 2kb of flanking. Three sequences are mapped: full insertion with flanking (“full”), left and right flanking only (“left” and “right”). The predicted insertion sequence from the genome GIM-012 fully maps onto the reference dm6, with only an additional 6bp insertion in 5’ (orange triangle).

**2R_21826762_21826762_roo**
Expected TE: roo
Expected SV length: 8045 bp
Expected in sample: RAL-091

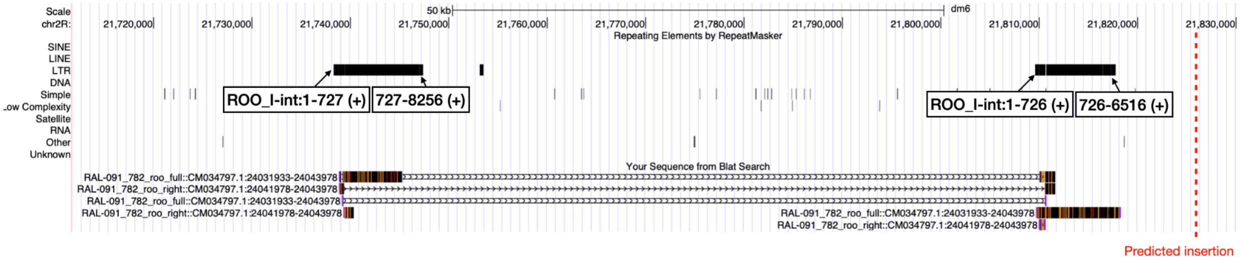


**Supplementary Figure 7. False positive insertion in the dm6 assembly reported by Rech et al., 2022.** Screenshot from the UCSC genome browser showing Blat mapping of the predicted insertion sequence +/- 2kb of flanking. Three sequences are mapped: full insertion with flanking ("full"), left and right flanking only ("left" and "right"). The predicted insertion maps ambiguously onto the reference dm6, attracted by two existing roo insertions located 50kb apart. There is no evidence that the flanking sequences of the predicted insertion maps in the non-repetitive region where the insertion is predicted (red dashed bar).

**2R_6482661_6482661_Micropia**
Expected TE: Micropia
Expected SV length: 338 bp
Expected in sample: SLA-001

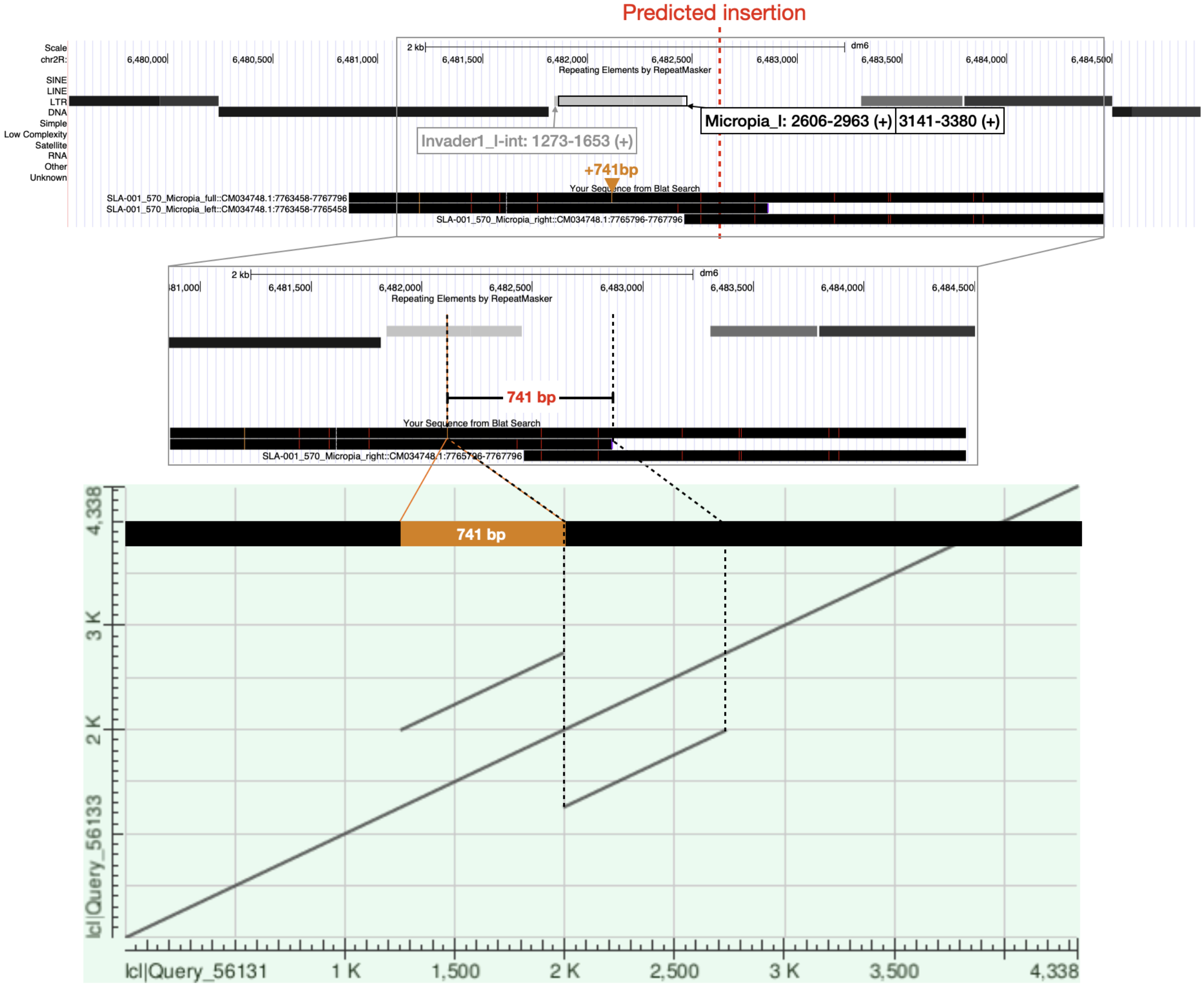


**Supplementary Figure 8. False positive insertion in the dm6 assembly reported by Rech et al., 2022.** Screenshot from the UCSC genome browser showing Blat mapping of the predicted insertion sequence +/- 2kb of flanking. Three sequences are mapped: full insertion with flanking ("full"), left and right flanking only ("left" and "right"). The mapping indeed provides evidence for an 741 bp tandem duplication (shown in the self dot-plot of the "full" sequence containing the predicted TE insertion and +/- 2kb of flanking, which includes 333 bp of a Micropia element, previously annotated as Invader1 on the reference genome.

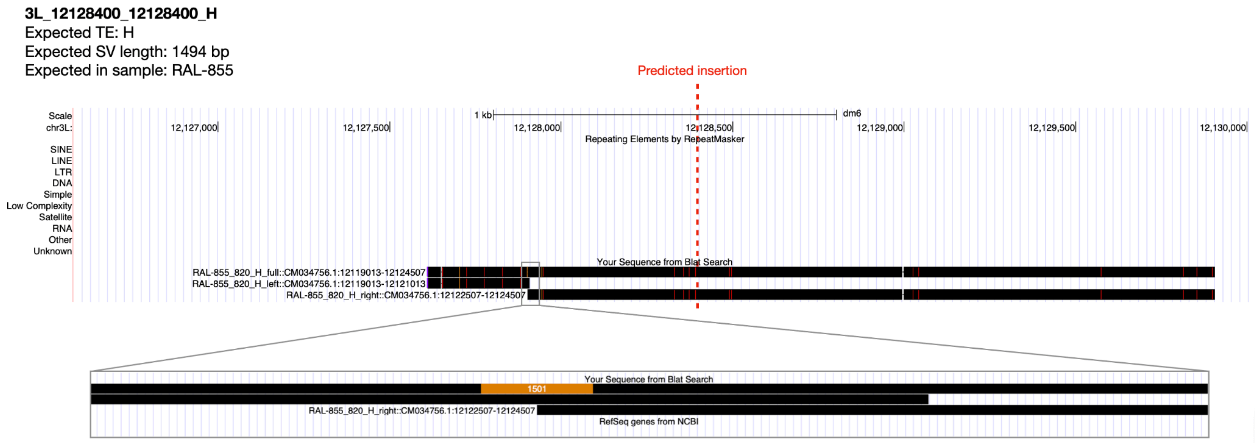


**Supplementary Figure 9. True positive insertion in the dm6 assembly reported by Rech et al., 2022.** Screenshot from the UCSC genome browser showing Blat mapping of the predicted insertion sequence +/- 2kb of flanking. Three sequences are mapped: full insertion with flanking ("full"), left and right flanking only ("left" and "right"). Note the distance of ~500bp between the predicted (red dashed line) and actual (orange strip) breakpoint.

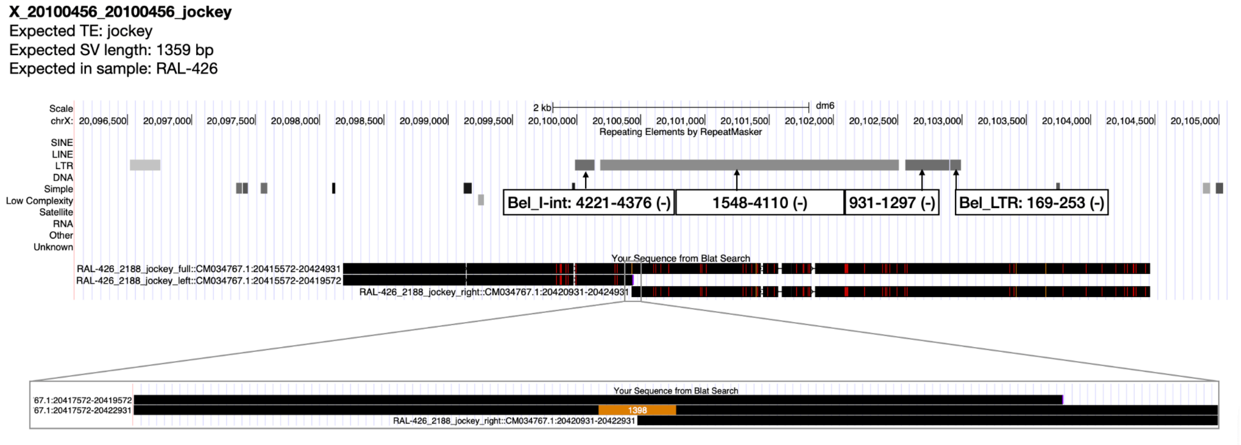


**Supplementary Figure 10. True positive insertion in the dm6 assembly reported by Rech et al., 2022.** Screenshot from the UCSC genome browser showing Blat mapping of the predicted insertion sequence +/- 4kb of flanking (flanking was extended 2kb to anchor them outside of the Bel elements present in the reference genome). Three sequences are mapped: full insertion with flanking (“full”), left and right flanking only (“left” and “right”). This insertion is nested in a Bel element present in the reference genome dm6.

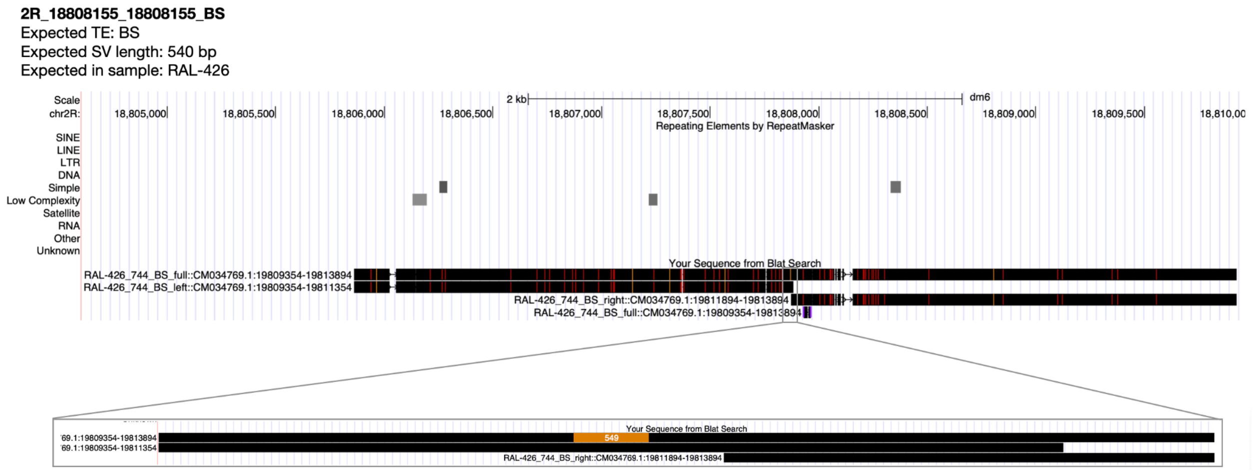


**Supplementary Figure 11. True positive insertion in the dm6 assembly reported by Rech et al., 2022.** Screenshot from the UCSC genome browser showing Blat mapping of the predicted insertion sequence +/- 2kb of flanking. Three sequences are mapped: full insertion with flanking ("full"), left and right flanking only ("left" and "right").

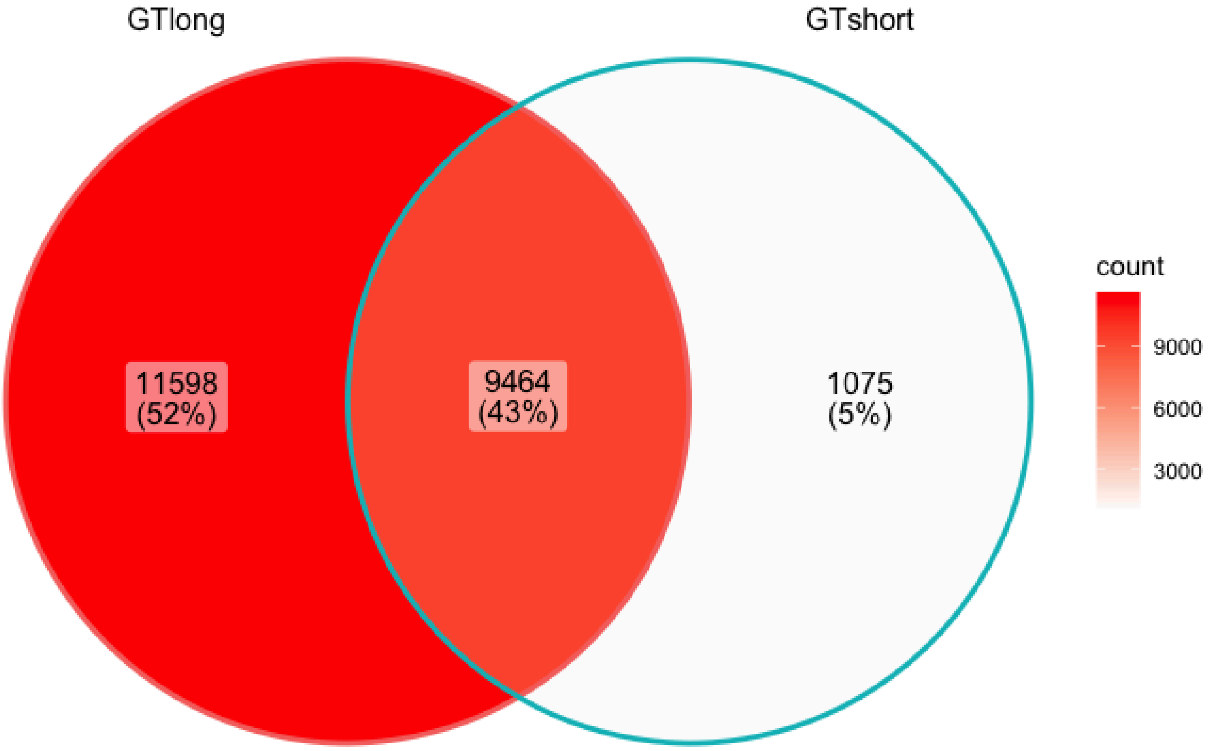


**Supplementary Figure 12. Comparison of non-fixed variants counts between GT-svsn-GA mode (long read genotyping) and GT-svsn-PG (short read genotyping).** For each dataset (VCF file) only variants with a single hit to a TE consensus were kept. Fixed variants post-genotyping and variants with missing genotype were also removed.

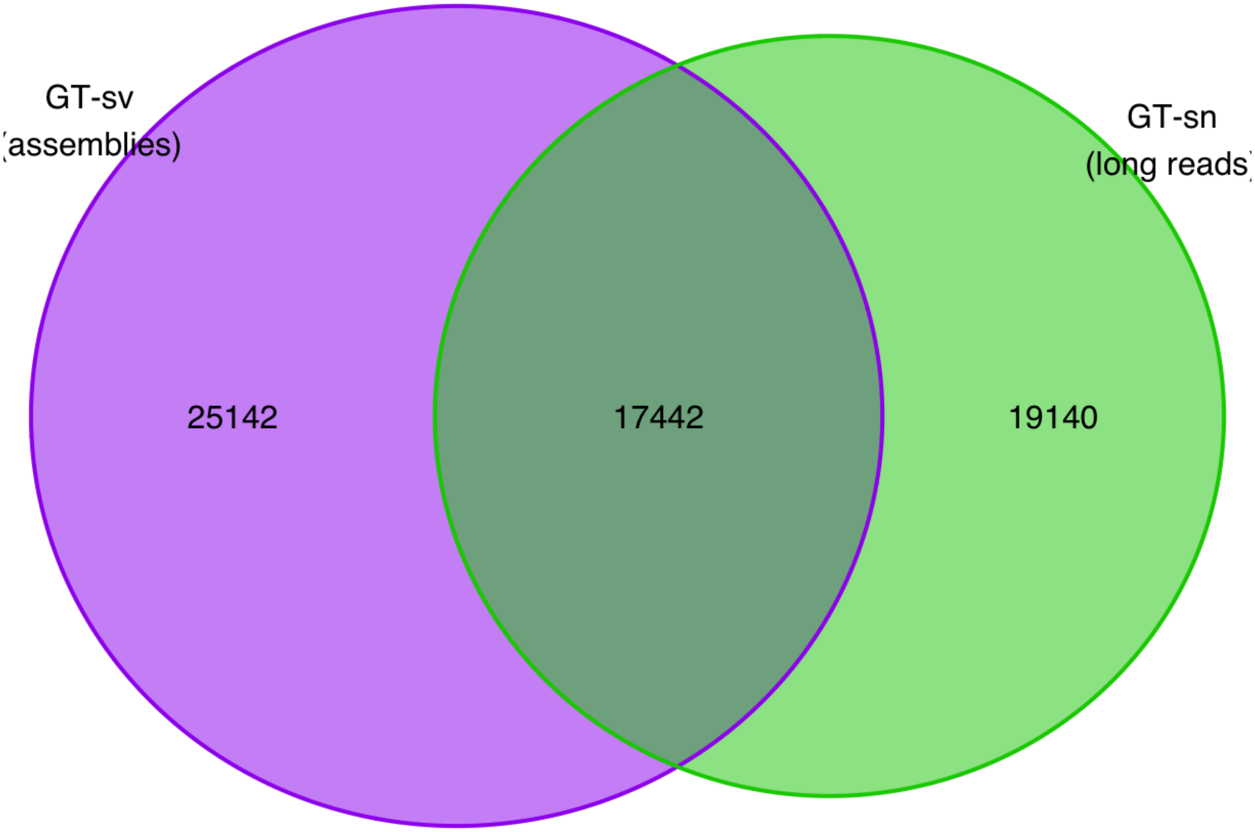


**Supplementary Figure 13. Comparison of the total number of candidate pME in the *C. sativa* pangenome whether assemblies (GT-sv) or long-reads (GT-sn) are used for pME search.** Search from long-read is restricted to 5 samples for which long-reads were publicly available, while GT-sv was run on 9 assemblies. Note that the total number of pMEs retained in GT-sv mode is slightly higher (42584 *vs.* 42517) than in the main text: indeed, clustering of candidate loci, as described in the manuscript, is performed on the total set (GT-sv + GT-sn) and clusters of 3+ loci are retained. Thus, more loci (provided by the GT-sn workflow) are available to reach clusters sizes of 3 and be retained in the analysis. Using long-reads adds a total of 19,207 loci, including 67 from GT-sv now present in clusters of 3+ loci.

### Supplementary Tables

The supplementary tables are available as separate Excel spreadsheets.

**Supplementary Table 1 A:** Combinations of SV search and genotyping coverages performed for GraffiTE and other methods evaluation. **B:** Methods' acronyms legend

**Supplementary Table 2:** Manual comparison between Rech et al., 2022 pME predictions and GraffiTE output for 94 insertions predicted by Rech et al. and not found in GraffiTE results using the sveval package

**Supplementary Table 3.** Information relative to *Cannabis sativa* genome assemblies used with GraffiTE

**Supplementary Table 4.** Computational resources usage report. Examples of CPU (available), RAM (peak usage) and runtime for diverse GraffiTE processes. RAM is rounded up to the next Gb, time is rounded up to the next minute.

**Supplementary Table 5.** Dot-plot analysis of the structural variants containing LTR/Copia annotations at the *bz* locus of *Zea mays*
